# Pathogenic neutrophilia drives acute respiratory distress syndrome in severe COVID-19 patients

**DOI:** 10.1101/2021.06.02.446468

**Authors:** Devon J. Eddins, Junkai Yang, Astrid Kosters, Vincent D. Giacalone, Ximo Pechuan, Joshua D. Chandler, Jinyoung Eum, Benjamin R. Babcock, Brian S. Dobosh, Mindy R. Hernández, Fathma Abdulkhader, Genoah L. Collins, Richard P. Ramonell, Christine Moussion, Darya Y. Orlova, Ignacio Sanz, F. Eun-Hyung Lee, Rabindra M. Tirouvanziam, Eliver E.B. Ghosn

## Abstract

Severe acute respiratory syndrome coronavirus 2 (SARS-CoV-2) and the ensuing COVID-19 pandemic have caused ∼40 million cases and over 648,000 deaths in the United States alone. Troubling disparities in COVID-19-associated mortality emerged early, with nearly 70% of deaths confined to Black/African-American (AA) patients in some areas, yet targeted studies within this demographic are scant. Multi-omics single-cell analyses of immune profiles from airways and matching blood samples of Black/AA patients revealed low viral load, yet pronounced and persistent pulmonary neutrophilia with advanced features of cytokine release syndrome and acute respiratory distress syndrome (ARDS), including exacerbated production of IL-8, IL-1β, IL-6, and CCL3/4 along with elevated levels of neutrophil elastase and myeloperoxidase. Circulating S100A12^+^/IFITM2^+^ mature neutrophils are recruited via the IL-8/CXCR2 axis, which emerges as a potential therapeutic target to reduce pathogenic neutrophilia and constrain ARDS in severe COVID-19.

**Graphical Abstract:** The lung pathology due to severe COVID-19 is marked by a perpetual pathogenic neutrophilia, leading to acute respiratory distress syndrome (ARDS) even in the absence of viral burden. Circulating mature neutrophils are recruited to the airways via IL-8 (CXCL8)/CXCR2 chemotaxis. Recently migrated neutrophils further differentiate into a transcriptionally active and hyperinflammatory state, with an exacerbated expression of IL-8 (*CXCL8*), IL-1β (*IL1B*), *CCL3, CCL4*, neutrophil elastase (NE), and myeloperoxidase (MPO) activity. Airway neutrophils and recruited inflammatory monocytes further increase their production of IL-8 (*CXCL8*), perpetuating lung neutrophilia in a feedforward loop. MdCs and T cells produce IL-1β and TNF, driving neutrophils reprogramming and survival.

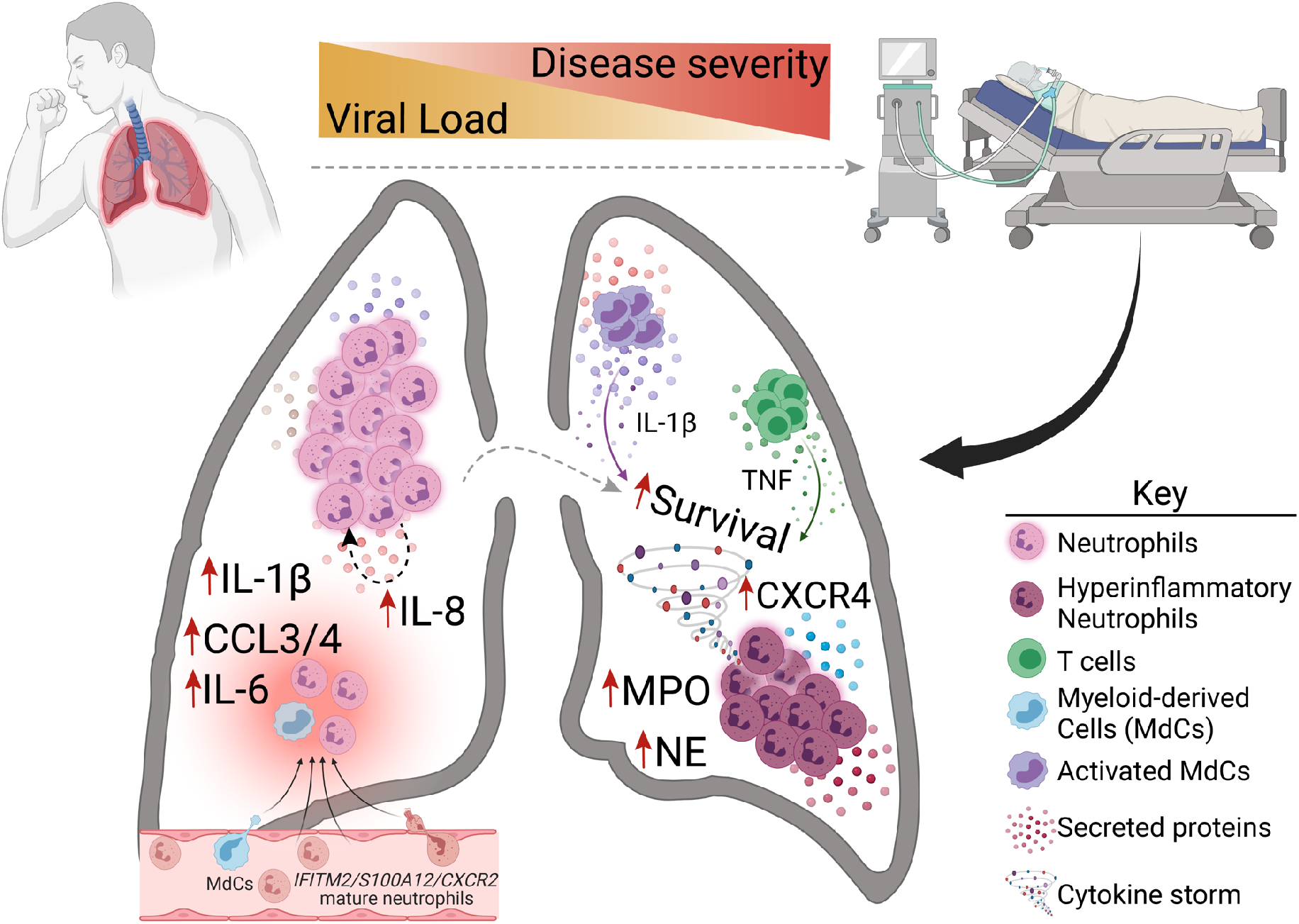

## Introduction

The coronavirus disease 2019 (COVID-19) pandemic, caused by SARS-CoV-2, is associated with high morbidity and mortality. To date, the pandemic has caused ∼40 million cases and over 648,000 deaths in the United States alone, where communities of color are disproportionately burdened with disease severity and mortality^1, 2^. A hallmark of COVID-19 pathogenesis is the vast array of clinical presentations and outcomes, ranging from asymptomatic or mild, self-limiting disease to acute respiratory distress syndrome (ARDS), multiorgan failure, and death. Indeed, such diversity in COVID-19 pathogenesis poses challenges to identify processes that dictate progression to severe disease. Early reports highlighted systemic hyperinflammatory responses that are linked to disease severity^3^. Cytokine release syndrome, also called a cytokine storm, has been observed in many patients and is suspected of causing the detrimental progression of COVID-19 and sustained immune dysregulation^4^.

Severe COVID-19 parallels the pathophysiology of sepsis^5^, where clinical presentation often includes granulocytosis, elevated proinflammatory cytokine production, aberrant myeloid activation, altered dendritic cell (DC) population dynamics, and lymphopenia^6, 7^. Early single-cell analyses of bronchoalveolar lavage fluid (BALF) samples implicated dysregulated monocyte and macrophage responses as central features in poor outcomes^4, 8^. As such, most early efforts have been focused on characterizing and constraining aberrant monocyte/macrophage responses. Concurrently, reports of neutrophilia in the peripheral blood arose, and the neutrophil to lymphocyte ratio emerged as an independent risk factor for disease progression^9^. Interestingly, a prior investigation of ARDS following sepsis identified sustained neutrophilia associated with worsened prognosis and death compared to patients who resolved neutrophilia and showed an increase in tissue-resident alveolar macrophages^10^, which may present another shared feature between COVID-19 and sepsis.

To date, there have been many studies of neutrophil responses in the blood of COVID-19 patients^11, 12, 13, 14, 15^. However, an in-depth analysis of the neutrophil activity in the lungs is lacking. Expressly, the extent to which neutrophils contribute to cytokine release syndrome, tissue damage, and ultimately ADRS in severe COVID-19 cases is incompletely understood. Our systems immunology approach, combining high-dimensional (Hi-D, 30-parameter) flow cytometry and multi-omics single-cell sequencing analyses of immune profiles from the airways and matching blood samples of Black/African-American (AA) patients, revealed pronounced and sustained pulmonary neutrophilia as a hallmark of severe disease. Furthermore, mature pulmonary neutrophils produce very high levels of the neutrophil chemotactic factor IL-8 in addition to IL-1β, IL-6, and CCL3/4 along with copious amounts of neutrophil elastase (NE) and myeloperoxidase (MPO). Altogether, our findings highlight that transcriptionally active and highly inflammatory neutrophils are sustained in the airways of severe patients and that reducing pathogenic neutrophilia may constrain ARDS in severe COVID-19 disease.

## Results

### Study Cohort

To better understand immune dynamics in Black/AA patients with severe COVID-19, we analyzed airway and matching blood samples from a cohort of 35 individuals presenting to Emory University Hospitals (severe) or the Emory Acute Respiratory Clinic (mild-acute) in Atlanta, GA, USA (Fig. 1 and Extended Data Table 1), including 8 demographic-matched healthy adults as controls. Of the 27 individuals who were confirmed positive by PCR from nasopharyngeal swabs, 18 had an NIH severity score of “critical” (https://www.covid19treatmentguidelines.nih.gov/overview/clinical-spectrum/; referred to as severe herein) and were admitted to the intensive care unit (ICU) requiring mechanical ventilator support, and 9 were mild (mild-acute) outpatients. All severe patients in our cohort received corticosteroids (dexamethasone or equivalent). Approximately half received one or more doses of the antiviral medication remdesivir, with an average ICU stay of 26 days (see Extended Data Tables 1 and 2).

**Figure 1.**
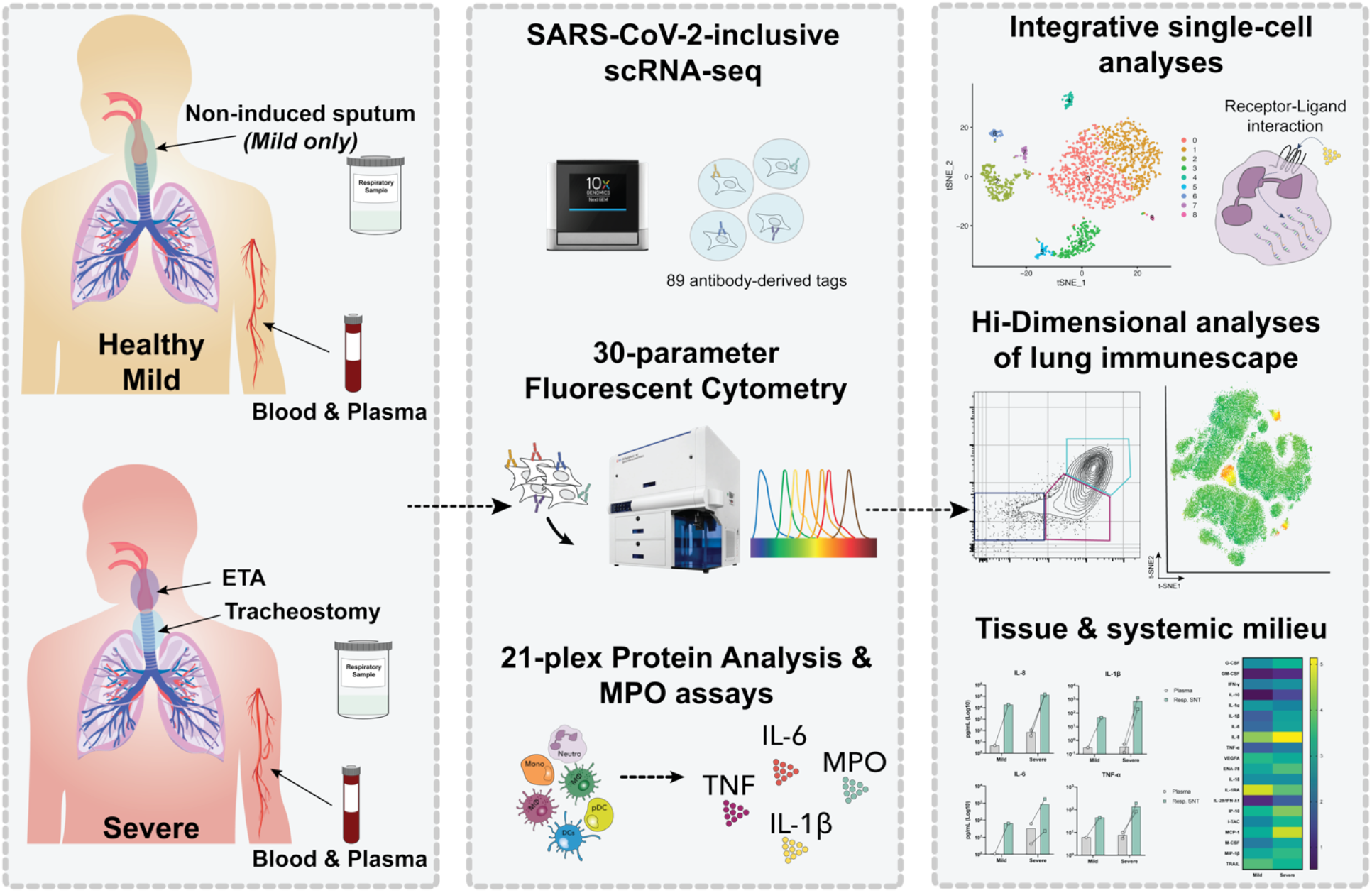
Experimental design for the systems immunology approach (integrated multi-omics single-cell assays) to study COVID-19 patient samples. Respiratory samples (sputum or endotracheal aspirates) and matching blood from all subjects were collected for analysis by 21-plex Mesoscale analysis, high-dimensional (Hi-D) 30-parameter flow cytometry, and multi-omics scRNA-seq. Cells from endotracheal aspirates (ETA) and blood of severe COVID-19 patients along with blood from healthy individuals were surface-stained with a panel of 89 oligo-conjugated monoclonal antibodies before single-cell encapsulation, and analyses were performed with a custom human reference genome that included the SARS-CoV-2 genome to simultaneously detect viral mRNA transcripts. Integrative multi-omics analyses were performed on the resulting data sets.

### Exacerbated neutrophilia in the airways and matching blood of severe COVID-19 patients

We first characterized major immune lineages in the airways (endotracheal aspiration, ETA) and matching blood samples by Hi-D flow cytometry and observed a pronounced circulating neutrophilia and lymphopenia (notably T and NK cells), which is similarly reflected in the airways (Fig. 2a-e and Extended Data Fig. 1). This is in line with recent reports showing lymphopenia associated with significant alterations in the myeloid compartment^15, 16^. Strikingly, in most cases of severe disease, ≥85% of all pulmonary leukocytes were neutrophils (Fig. 2b). This contrasts with other studies that have reported much more heterogeneity in neutrophil frequency in patients’ blood and lungs^17, 18^. In addition, circulating T cells and NK cells decline with increased disease severity (Fig. 2d), which has recently been reported as a feature of COVID-19 pneumonia^19^. We also observed a notable decrease in pulmonary NK cells associated with disease severity (Fig. 2b). However, there were only subtle differences in the B-cell compartment (which were detected at very low levels in the airways) and myeloid-derived cells (MdCs) compartment compared to other reports^8, 20, 21^, highlighting the importance of neutrophilia and neutrophil-to-lymphocyte ratio in our cohort. Interestingly, the patient with the lowest neutrophil frequency in the airways corresponds to the youngest patient in the severe disease group (Fig. 2b). However, since we only had a single patient under 35 years old in our severe group, we cannot conclude any associations between neutrophilia and patient age.

**Figure 2.**
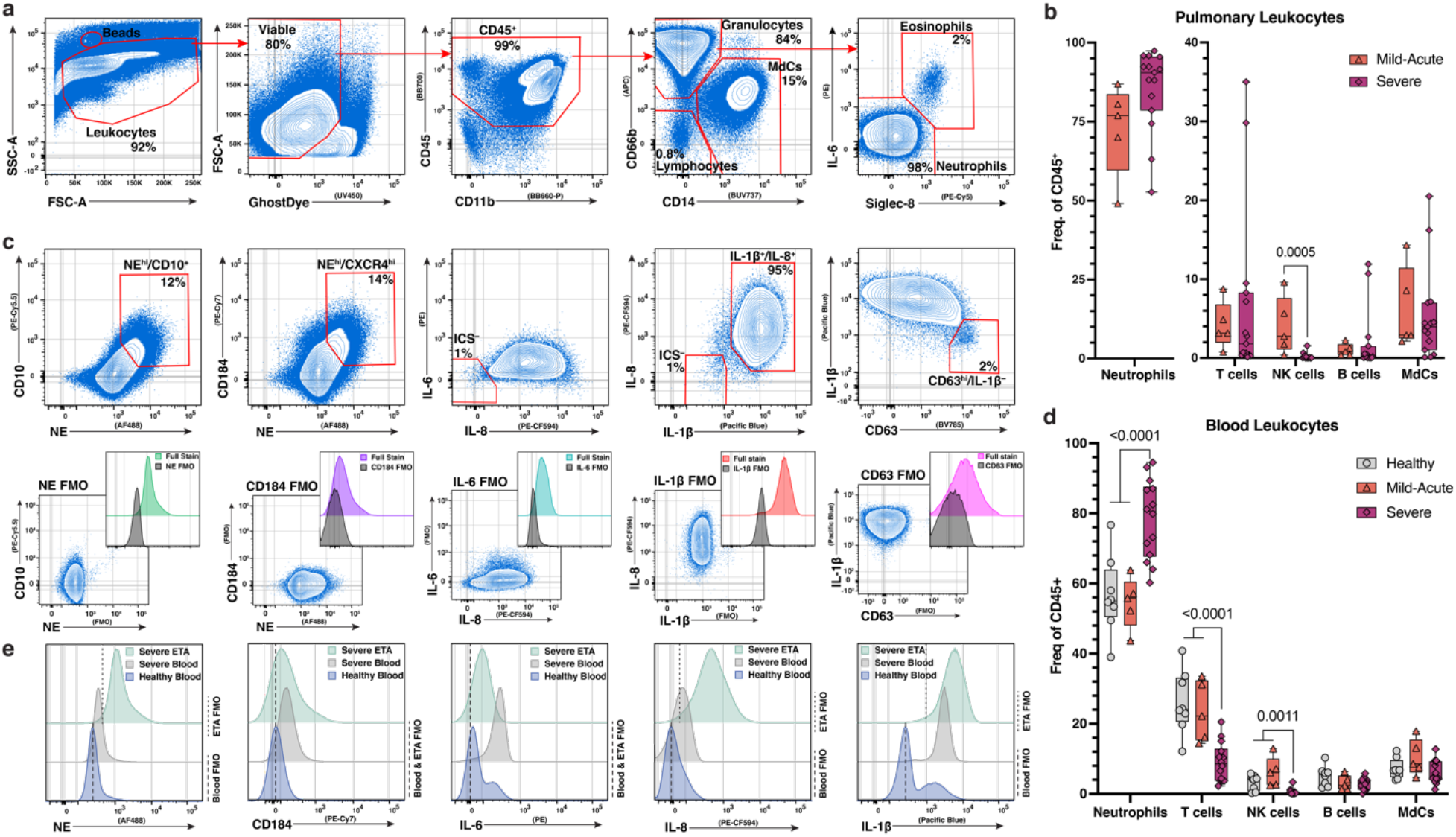
Exacerbated neutrophilia in the airways and matching blood of severe COVID-19 patients. (a) Representative gating strategy for all samples (see Extended Data Fig. 1 for full gating strategy). (b) Box plots show distributions of leukocytes isolated from endotracheal aspirates (ETA). (c) Representative plots demonstrating inflammatory profile of pulmonary neutrophils including neutrophil elastase (NE), CD184 (CXCR4), and intracellular staining of IL-6, IL-8, and IL-1β including the full stain and fluorescence minus one (FMO) controls (d) Box plots show distributions of leukocytes isolated from whole blood from severe patients. (e) Representative histograms showing median fluorescence intensity (MFI) of key markers across healthy blood (blue), severe blood (gray), and severe ETA (green) samples. MdCs = myeloid-derived cells. For comparisons across the three patient groups (i.e., healthy, mild-acute, severe), ordinary one-way ANOVA (if equal variance) or Brown-Forsythe and Welch ANOVA (if unequal variance) tests were performed for data with a normal distribution. Data with a lognormal distribution were analyzed with a Kruskal-Wallis test.

Since we identified prominent neutrophilia in our cohort, we developed a Hi-D, 30-parameter flow cytometry panel to interrogate inflammatory neutrophil phenotype in addition to broad leukocyte characterization (see “CoV-Neutrophil” panel in Extended Data Table 3). Here, we used extracellular staining of neutrophil elastase (NE) and CD63 (LAMP3) to assess primary granule release^22^, coupled with intracellular staining for key effectors implicated in cytokine storm (IL-1β, IL-6, and IL-8). We observed that nearly all neutrophils recruited to the lung developed an inflammatory profile characterized by exacerbated levels of NE and elevated production of IL-1β and IL-8 with a dramatic increase in primary granule release (Fig. 2c,e). Indeed, most neutrophils in the airways were positive for all 3 intracellular cytokines and express CD63 on the cell surface (Fig. 2c), suggesting that neutrophils are releasing cytokines upon degranulation. However, this phenotype is diminished in the circulation. A smaller proportion of blood neutrophils undergo granule release and express a cytokine signature limited mainly to IL-1β production and less IL-6 and IL-8 (Fig. 2e). Similarly, we find a subset of neutrophils in the lung that co-express the highest levels of surface CD10, CD184 (CXCR4) and NE (Fig. 2c). Interestingly, neutrophils in the lung express lower levels of FcγRII (CD32) than those in the blood (Extended Data Fig. 5a). Together, these results demonstrate systemic neutrophilia in severe COVID-19 patients where neutrophils accumulating in the airways produce exacerbated levels of potentially damaging enzymes (e.g., NE) and inflammatory cytokines (IL-1β/6/8) while undergoing pronounced primary granule release.

### Neutrophil-secreted inflammatory molecules in the airways potentiate acute respiratory distress syndrome in severe patients

To better understand the inflammatory signaling milieu and relate the intracellular cytokine staining to protein secretion, we measured 21 total analytes in the airways and plasma using the Mesoscale UPLEX platform (Extended Data Table 4). We found that IL-8 is the most abundant chemokine in the airways of severe patients accompanied by high levels of IL-1β and IL-6. This is in line with our Hi-D flow cytometry data showing neutrophilia and pronounced intracellular staining signal for IL-8 and IL-1β (Fig. 3a-i). Strikingly, IL-8 levels were ∼100-fold higher in the airways of severe versus mild patients and that of IL-1β and IL-6 levels in severe patients. These findings are in stark contrast to an early report^23^ that showed no differences between circulating and pulmonary IL-6 and IL-8 secretion, reported in the <100 pg/mL range. However, more recent studies in both ETA^24^ and BALF^25^ samples have noted similar findings that corroborate our own. These data, therefore, implicate IL-8 signaling in the prominent neutrophilia observed in severe patients in our cohort.

**Figure 3.**
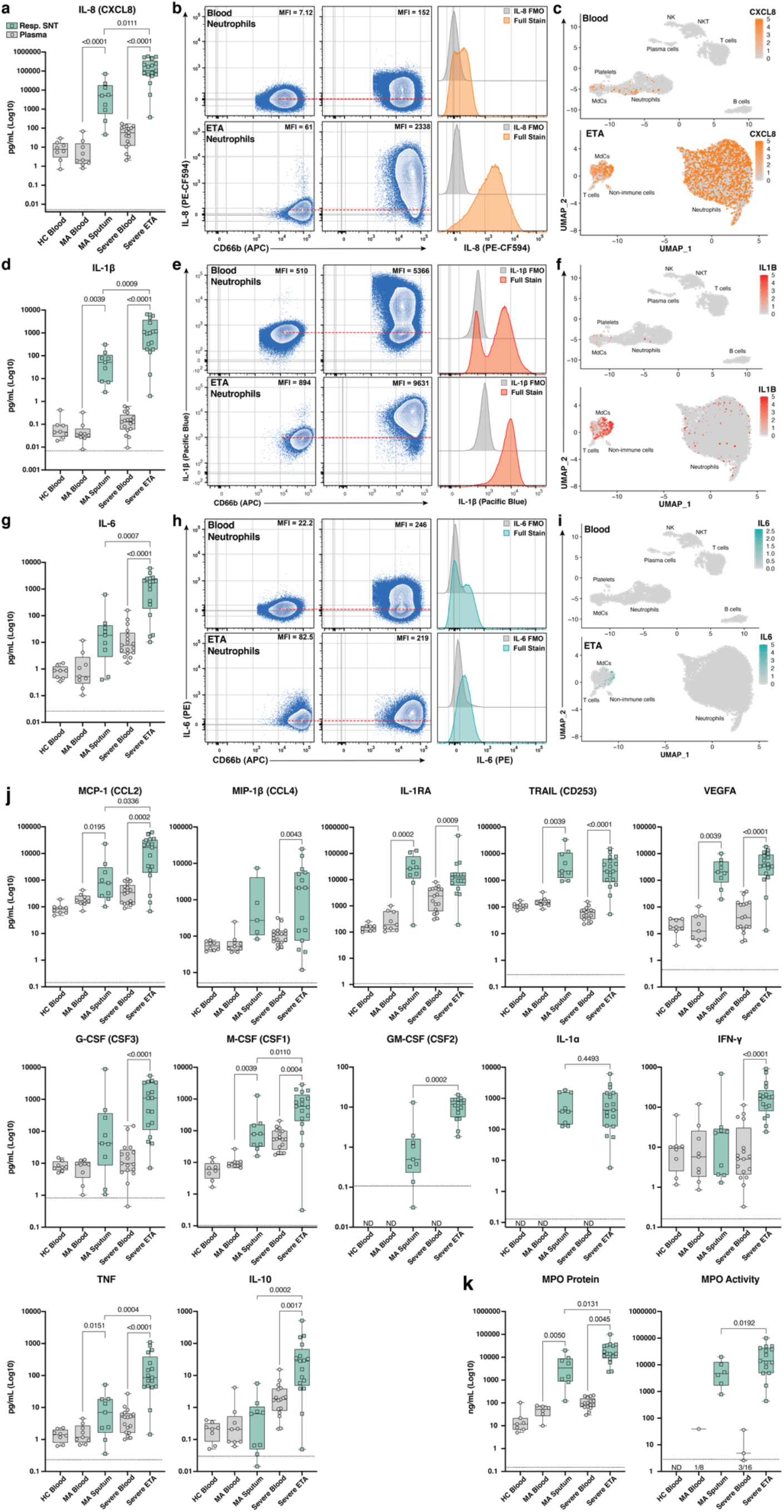
Cytokine release syndrome is dominated by IL-8 and IL-1β with pronounced myeloperoxidase content and activity in the lung microenvironment. (a, d, g, j) Concentration (pg/mL) of 15 analytes interrogated by Mesoscale analyses in plasma (gray circles) and respiratory supernatant (Resp. SNT; green squares) from healthy control (HC), mild-acute (MA), and severe COVID-19 patients. (b, e, h) Representative flow cytometric intracellular staining for IL-8 (CXCL8), IL-1β, and IL-6, including the full stain and fluorescence minus one (FMO) controls in both blood and ETA neutrophils (CD66b^+^). (c, f, i) UMAP visualizations of *CXCL8* (IL-8), *IL1B*, and *IL6*, which were also measured by intracellular flow cytometry staining and Mesoscale in both blood and ETA. (k) Concentration (ng/mL) of myeloperoxidase (MPO) protein and MPO activity in plasma vs. Resp. SNT. In (a, d, g, j, k), black dotted line = median lower limit of detection (LLOD) for assays (see Extended Data Table 3). In (b, e, h) red dashed line indicates the median fluorescence intensity (MFI) of neutrophils in the FMO control (value listed in the plot).

We noticed only minor differences in circulating levels of many targets between healthy individuals and those affected by severe COVID-19 disease suggesting an uncoupling of local versus systemic inflammatory responses^24^, particularly for IL-8 and IL-1β, but also IL-6 to a lesser extent (Fig. 3a,d,g). Conversely, we observed a rise in circulating IL-1RA levels with increasing disease severity, similar to other reports^26^. However, we observed an opposite trend in the airways, where IL-1RA levels appear to be decreased in severe patients (Fig. 3j). This is in stark contrast to IL-1β levels, which were not markedly different in the circulation but significantly increased in the airways with disease severity (Fig. 3j). We also found increased CCL2 (MCP-1) in the airways of severe disease, and the same trend for M-CSF and TNF was observed, albeit at much lower concentrations (Fig. 3j). Interferon (IFN)-γ was detected at the highest levels in the airways of severe patients with no obvious differentiation in plasma across all groups. IL-10 was also increased in the airways of severe patients with notable variation at low levels (Fig. 3j). In contrast to other studies^27, 28^, we did not observe appreciable differences in IL-18 or CXCL10 (IP-10) levels across patient groups (Extended Data Fig. 2). Additionally, CXCL12 was not detected in the airways of patients, and there were no notable differences in plasma levels across patient groups (Extended Data Fig. 2).

We further measured the concentration of neutrophil-derived MPO and its enzymatic activity across patient groups. MPO is an important neutrophil effector molecule (originating from primary granules like NE) implicated in respiratory illnesses such as cystic fibrosis^29^, and can contribute to lung tissue damage during neutrophilic pneumonitis. MPO protein concentrations increased stepwise with disease severity in the plasma and respiratory supernatant (Fig. 3k). However, MPO activity was mostly detected in the airways, and rose with increasing disease severity (Fig. 3k).

### Recruited airway neutrophils are mature, transcriptionally active, and further differentiate into a highly inflammatory state

To gain insight into the cellular states and transcriptional regulation in patients with severe COVID-19 disease, we performed multi-omics scRNA-seq on cells from whole blood and ETA samples of severe patients and whole blood from demographic-matched healthy control (Fig. 1). We assessed immune features in data integrated from four or more patients from the same cohort (i.e., healthy vs. severe; see Methods). First, major lineages from blood (Fig. 4a) and ETA (Fig. 4g) were gated manually using the antibody-derived tag (ADT) data for surface protein expression. Next, we used the gene expression (GEX) data to generate clusters for each major lineage identified based on their surface markers, as per our previously described SuPERR-seq pipeline^30^ (Extended Data Fig. 3). For example, neutrophils identified by CD66b and CD16 surface ADT in the blood (Fig. 4b) and ETA (Fig. 4h) were then selected and clustered independently by GEX data (Fig. 4c,i) for further analyses.

**Figure 4.**
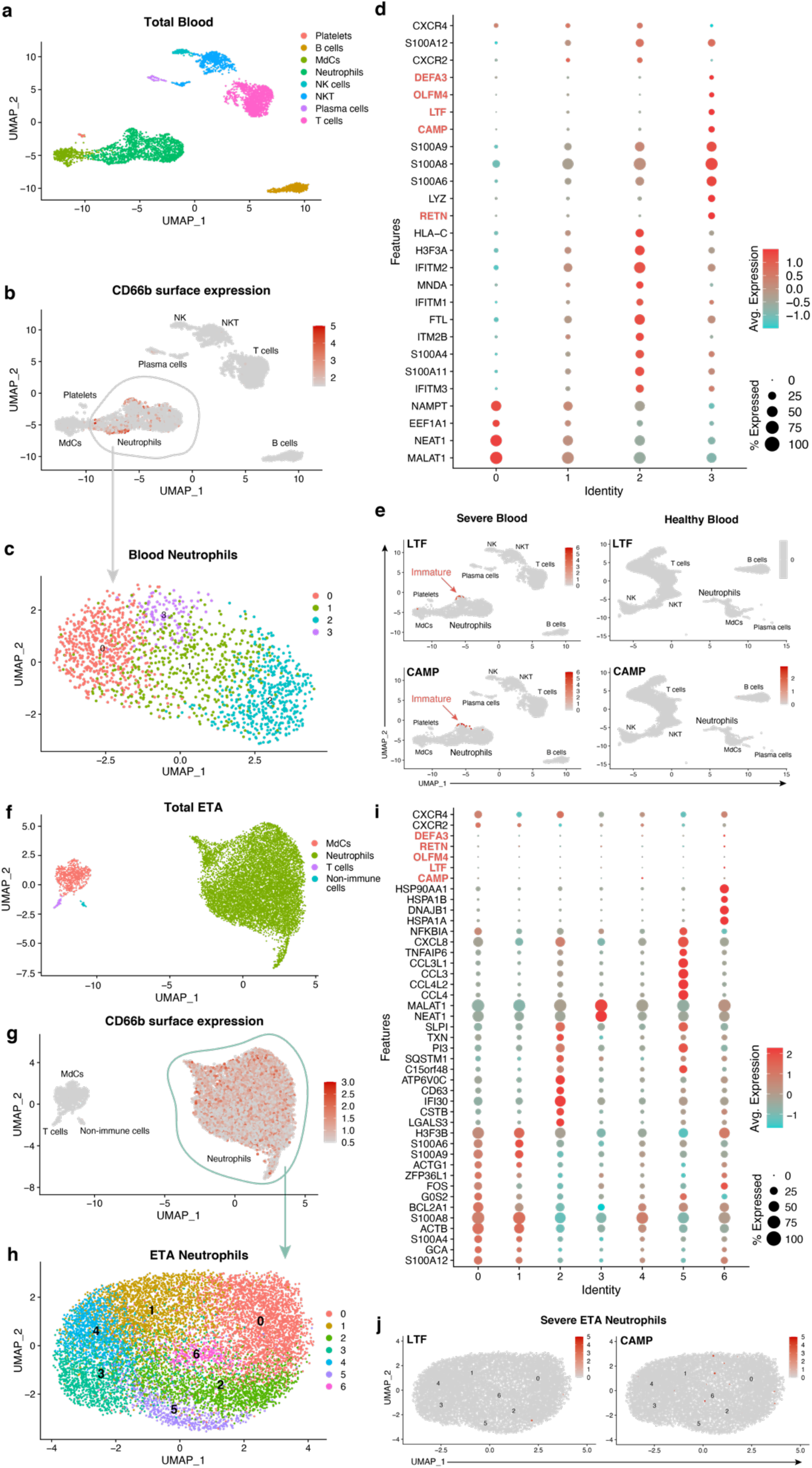
Multi-omic single-cell RNA-seq reveals emergency granulopoiesis in the circulation and abundant heterogeneous populations of mature neutrophils in the airways with distinct inflammatory states. UMAP visualization of scRNA-seq of total integrated blood (a) and endotracheal aspirate (ETA) (f) cells. Neutrophils were identified based on cell-surface markers (b and g), and total neutrophils were subclustered for further analysis (c and h). Dot plots of the intersection of the top differentially expressed genes in neutrophil clusters (d and i) sorted by average log-fold change for blood (d) and lung (i) neutrophils, respectively. UMAP visualization of signature genes of immature neutrophils (highlighted in d and i) in blood from severe patients compared to healthy individuals (e) and lungs of severe patients (j).

A prior study had reported an increase in immature neutrophils in the lung concomitant to increased circulating immature neutrophils from emergency hematopoiesis in severe COVID-19 patients^16^. We, therefore, assessed the immature neutrophil phenotype in the blood (Fig 4a-e) and lung (Fig 4f-j) by scRNA-seq in our patient cohort. We readily identified a cluster of neutrophils expressing *CAMP, LTF, RETN, OLFM4, DEFA3, CD24*, and *MMP8* (Fig. 4e and Extended Data Fig. 4) in the blood of severe patients (cluster 3 in blood neutrophils) that also has high expression of calprotectin (S100A8/9) and other calgranulins (Fig 4d). This is consistent with immature neutrophil phenotypes described by others^7, 15^ and was notably absent in neutrophils from healthy blood, confirming an emergency hematopoiesis/granulopoiesis in our patient cohort. However, in contrast to previous studies^16^, we did not observe any signature of immature neutrophils in the lungs of severe patients (Fig. 4j), suggesting that immature neutrophils either are not directly recruited to the lung, or quickly differentiate upon infiltration into the lung.

To determine which neutrophil subset in the blood can infiltrate the lung and further differentiate into a pathogenic state, we explored cell-cell communication (CellChat^31^) of clustered cells from blood (Fig. 4c) and lung (Fig. 4h) samples. We found significant communication through the CXCL pathway between the lung neutrophil, myeloid, and non-immune populations (i.e., epithelial/stromal cells) with *CXCR2*-expressing neutrophils from the blood (blood cluster 2; Fig. 5a). The CXCL8 (IL-8)/CXCR2 pathway was identified as the primary recruitment axis for circulating *CXCR2*^+^ neutrophils (Fig. 5b). These data are in line with our finding that IL-8 was increased in neutrophils by flow cytometry (Fig. 2) and secreted at very high levels in the airways (Fig. 3) and suggests that the CXCL8/CXCR2 signaling axis is important for neutrophil recruitment to the lungs during COVID-19 pathogenesis. Virtually all lung neutrophils (particularly cluster 5), non-immune cells, and monocytes show the potential to recruit circulating neutrophils (Fig. 5a,d), indicating a robust and redundant mechanism of neutrophil recruitment to the airways via the CXCL8/CXCR2 axis. Surprisingly, the immature neutrophils (blood cluster 3) lacked *CXCR2* (Fig. 5c), suggesting that immature neutrophils are unlikely to infiltrate the lung via IL-8. In contrast, a defined subset of mature neutrophils in blood expressing high levels of *CXCR2*, along with interferon-induced *IFITM2* and *S100A11/12*, identify blood cluster 2 as the putative neutrophil subset that can infiltrate the lung via IL-8 (Figs. 4d and 5c-d). It is, therefore, probable that recent lung emigrants would still express detectable levels of *CXCR2*, as well as *IFITM2* and *S100A11/12* (Fig. 5c).

**Figure 5.**
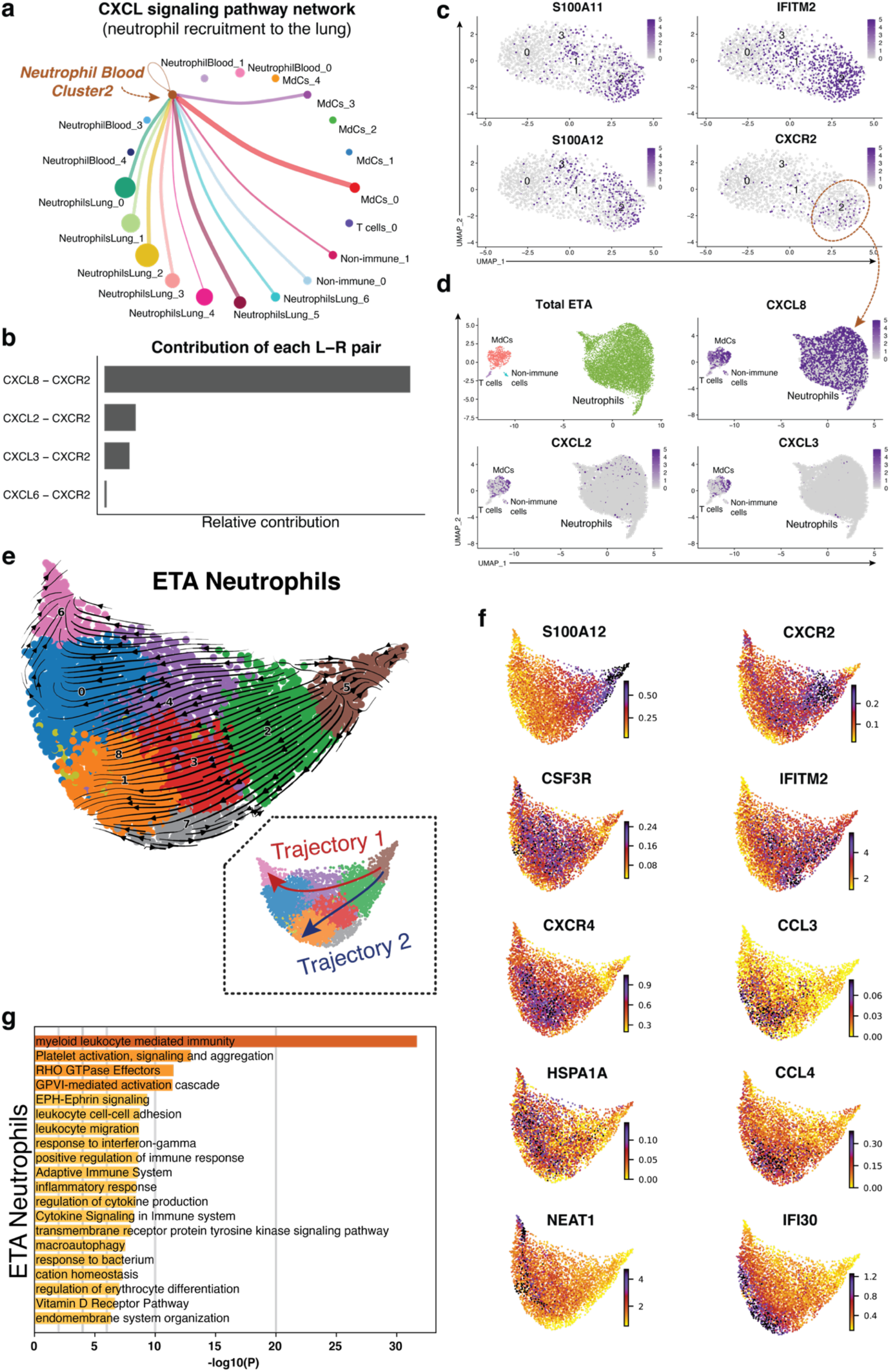
Mature neutrophils are continuously recruited from circulation and progress toward a hyperinflammatory state. Cell receptor-ligand pair analyses (CellChat^31^) from scRNA-seq data identify significant CXCL signaling pathway network enrichment between lung cells and blood neutrophils (a and b). Blood neutrophil cluster 2 represents the main subset potentially recruited to the lung (a). Recruitment of blood cluster 2 (a and c) is largely orchestrated by the CXCL8 (IL-8)/CXCR2 axis, and to a lesser extent, by the CXCL2 and CXCL3 (b and d). *S100A11/12, IFITM2*, and *CXCR2* mark a cluster of mature neutrophils in the blood (c) that likely represents the neutrophil subset recruited to the lung (d). Cell trajectory analysis (scVelo^32^) identifies two potential pathways (Trajectories 1 and 2) for recently migrated neutrophils (e), that begin with a gene signature consistent with neutrophil blood cluster 2 (c and f). Neutrophils recruited to the lung acquire a hyperinflammatory profile along Trajectory 2 (e and f), characterized by high expression of interferon-stimulated gene (ISG) *IFI30* along with macrophage inflammatory proteins *CCL3* (MIP-1α) and *CCL4* (MIP-1β), whereas *CSF3R* and *CXCR4* are increased in cells along both trajectories (f). Neutrophils along Trajectory 1 may reflect cells progressing to apoptosis expressing higher levels of *HSPA1A* (HSP70) followed by *NEAT1* (f). Pathway and process enrichment analyses performed in Metascape^85^ reveals that myeloid-mediated immunity and platelet activation, signaling, and aggregation are significantly enriched in neutrophils in the lungs of severe patients (g). MdCs: Myeloid-derived Cells.

After identifying the blood neutrophil cluster 2 as the putative source of lung-recruited neutrophils, we sought to identify their cell trajectory/differentiation once they enter the inflamed airway. Cell trajectory analyses (scVelo^32^) revealed two differentiation pathways (Trajectories 1 and 2) for the recently infiltrated neutrophils (Fig. 5e). Indeed, cells expressing the highest levels of *CXCR2* and *S100A11/12* were at the beginning of the trajectory, further supporting these as the putative emigrant population from the blood (Fig. 5f). Notably, neutrophils recruited to the lung and differentiated along Trajectory 2 experience transcriptional reprogramming to acquire a heightened inflammatory phenotype (Fig. 5e,f). In contrast to the canonical neutrophil differentiation pathway, including a short half-life, the infiltrated neutrophils in severe COVID-19 patients are transcriptionally active (Fig. 5e-g) in comparison to blood neutrophils (Extended Data Fig. 5b) and further differentiate into a hyperinflammatory state (Fig. 5f). This is consistent with more recent reports of neutrophil transmigration in other respiratory illnesses such as cystic fibrosis^33^, where neutrophils undergo lung/condition-specific adaptations upon recruitment from the circulation^34^ instead of the canonical rapid and transient effector function proceeding cell death. Indeed, we observed increased expression of the interferon-stimulated gene (ISG) *IFI30* along with increased expression of *CCL3* (MIP-1α) and *CCL4* (MIP-1β) (Fig. 5f), which in turn can promote the recruitment of inflammatory monocytes to the lung and/or proinflammatory macrophage phenotypes in the airways.

Importantly, most neutrophils increase expression of *CXCL8* (IL-8) when recruited to the airways (Fig. 5d), perpetuating neutrophil recruitment to the lung. These findings further corroborate our Hi-D flow cytometry, gene expression, and secreted protein data analysis identifying IL-8 as the most abundant neutrophil-derived chemokine (and neutrophil chemotactic factor) in the airways of severe COVID-19 patients. Furthermore, most neutrophils increase the expression of *CXCR4* when recruited to the airways (Fig. 5f). Interestingly, we find that *CXCR4* expression, previously attributed to immature neutrophils^16^, is increased in mature lung-recruited neutrophils (Fig. 5f). This is consistent with *CXCR4* expression dynamics previously detected in non-COVID lung inflammation, including in cystic fibrosis^35^ and malaria^36^. Taken together, the progressive increase in *IFI30, CCL3/4*, along with abundant *de novo CXCL8* and *CXCR4* mRNA transcripts, reveal a transcriptionally active state in neutrophils that is poised to sustain a hyperinflammatory milieu in the lung of severe COVID-19 patients.

### Viral load in the respiratory tract does not correlate with disease severity

We then investigated whether the recruited neutrophils or other cell types (i.e., myeloid, lymphoid, and non-immune cells) in the airways of severe patients were infected with SARS-CoV-2. By including the SARS-CoV-2 genome sequence into our human reference transcriptome in the multi-omics single-cell analysis, we were able to assess viral mRNA (vRNA) transcripts at a single-cell level (Fig. 7a). Notably, we did not detect vRNA in any cell types in the blood or airways of severe patients (Fig. 7b,c). Further, SARS-CoV-2-specific RT-qPCR revealed that viral load was decreased in the upper airways of severe patients admitted to the ICU compared to mild-acute patients (Fig. 7d,e). However, we did note a gene signature in neutrophils associated with response to IFN-γ (Fig. 5g and Extended Data Fig. 5b), including a pronounced increase in *IFITM2* (Extended Data Fig. 5c), which has been shown to promote SARS-CoV-2 infection in human lung cells^37^.

### Pulmonary TNF and IL-1β promote neutrophil reprogramming in the lungs

To ascertain which ligand-receptor pairs are potentially responsible for the transcriptional reprogramming of neutrophils in the lung, we first performed differential gene expression (DGE) analysis between blood neutrophil cluster 2 and lung neutrophils to identify genes that are upregulated in lung-recruited neutrophils (Fig. 6a). Next, we used the computational method NicheNet^38^ to identify the potential (prioritized) ligands that could induce the upregulation of the genes identified by the DGE analysis, indicating their potential role in neutrophil reprogramming in the lung. *TNF, IL1B*, and *APOE* were the highest prioritized ligands with high ligand activity whose signaling axes have the regulatory potential to drive gene expression profiles observed in lung-recruited neutrophils (Fig. 6b). Importantly, NicheNet analysis revealed *TNF* as the ligand predicted to increase *BCL2A1* expression in pulmonary neutrophils, as well as *CCL4* (MIP-1β) and *CXCL16*. Furthermore, both *TNF* and *IL1B* are the ligands predicted to induce *NFKBIA* and *CXCL8* (IL-8), and *IL1B* shows the greatest potential to induce *CCL3* (MIP-1α) expression in recruited neutrophils. *APOE* is predicted to upregulate the expression of *FCER1G* in neutrophils in the lung. Notably, *TNF, IL1B*, and *HMGB1* are the ligands that have the widest range of regulatory potential, inducing neutrophil reprogramming by upregulating most genes that we identified as differentially expressed (Fig. 6a) in lung-recruited neutrophils (Fig 6b).

**Figure 6.**
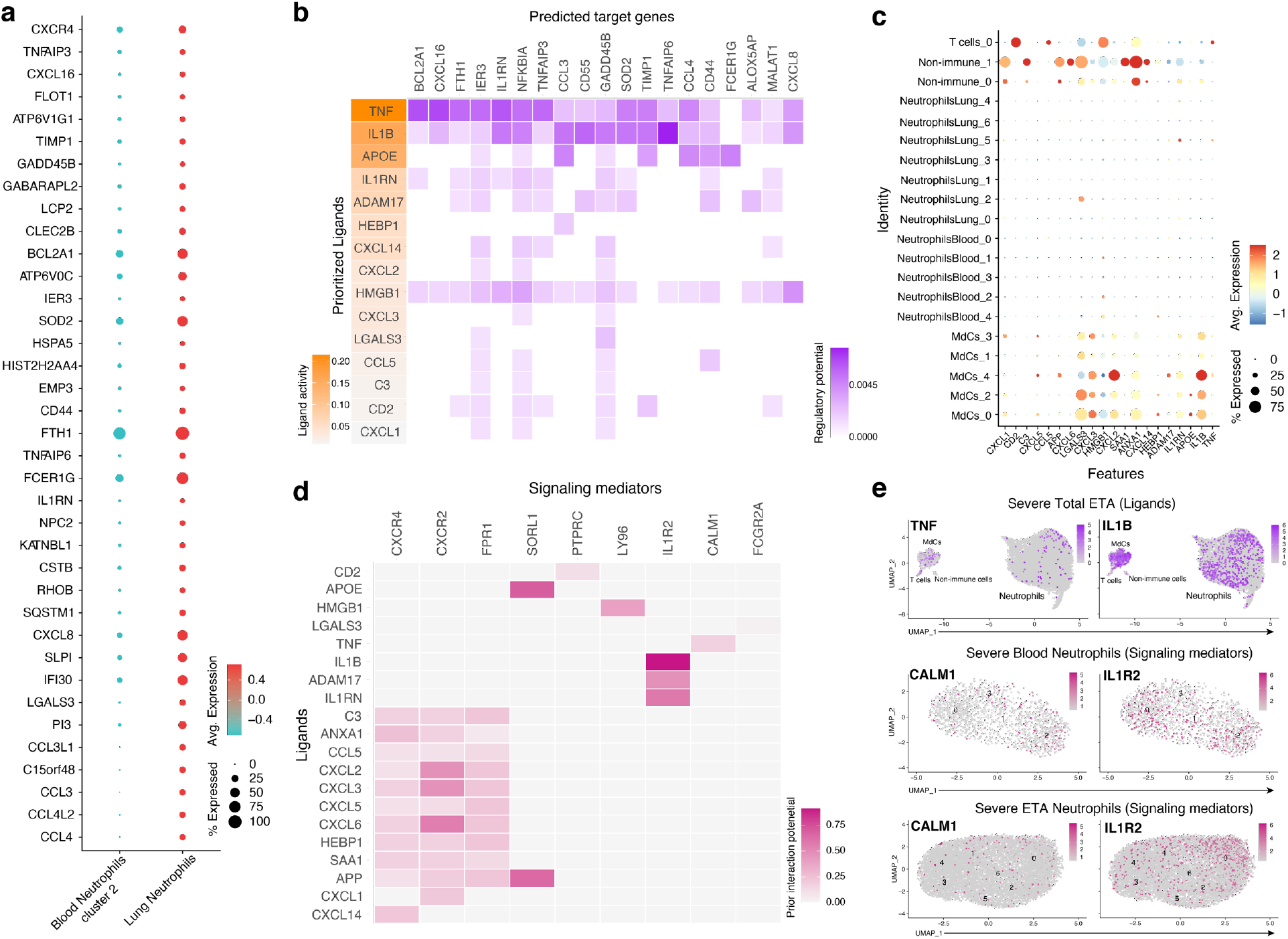
TNF and IL-1β drive inflammatory program in neutrophils recruited to the lung. Dot plots showing 37 of the 67 differentially expressed genes in lung neutrophils versus blood neutrophil cluster 2 sorted by average log-fold change used as input for NicheNet^38^ analyses (b) NicheNet^38^ analyses identified the highest prioritized ligands (top 15) ordered by ligand activity (y-axis) that best predict the pulmonary neutrophil gene signature (x-axis). The predicted target genes represent the pulmonary neutrophil gene signature identified by differential gene expression analysis between blood neutrophils from cluster 2 and ETA neutrophils (see Extended Data Fig. 5d) (b) Dot plots of the intersection of the top 15 expressed prioritized ligands from all cells in the ETA samples. (c) Ligand-receptor matrix of putative signaling mediators for the top 15 prioritized ligands identified in (a). (d) UMAP visualizations of *TNF* and *IL1B* transcripts in the total ETA along with expression of predicted signaling mediators in blood (middle) and ETA (bottom) neutrophils from severe patients.

Pulmonary T cells represent the primary cell population expressing *TNF* in the airways, where MdCs have the highest *IL1B* and *APOE* transcripts (Fig. 6c,e). From the transcriptome data, ligand-receptor pair analyses further identified putative signaling mediators for the top 15 prioritized ligands (Fig. 6d). *TNF* is predicted to signal through a receptor that has a *CALM1* association (Fig. 6d). Intriguingly, it has been previously reported that CALM1 can bind other transmembrane proteins, including ACE-2^39^, and regulate their cell surface expression^40^. *IL1B* is predicted to signal through the canonical IL1B/IL1R2 pathway, and *APOE* through the APOE/SORL1 lipid pathway. *HMGB1* is predicted to signal through *LY96* (MD-2), which is commonly associated with TLR4 (Fig. 6d)^41, 42, 43^. Accordingly, *IL1R2* and *CALM1* transcripts are abundant in neutrophils from blood cluster 2, and their expression are sustained upon recruitment to the airways (Fig. 6e).

Collectively, we show here that TNF- and IL-1β-mediated transcriptional reprogramming of lung infiltrating neutrophils leads to induction of proinflammatory *NFKBIA, CCL3* (MIP-1α), and *CCL4* (MIP-1β) along with *TNFAIP3/6* and *CXCL8* (Figs. 5 and 6). Of note, neutrophil-derived *CCL3* and *CCL4* can attract inflammatory monocytes from the circulation to the airways. Interestingly, lung-infiltrating monocytes also produce elevated levels of *CXCL8*, potentiating the recruitment of circulating neutrophils via the CXCL8/CXCR2 axis (Fig. 5a,d). Hence, both neutrophils and inflammatory monocytes in the airways exacerbate and potentiate pathogenic neutrophilia in the lungs of severe patients in a positive feedback loop.

## Discussion

Previous studies have reported increased neutrophilia in severe COVID-19, particularly in the circulation^14, 44, 45, 46^. In contrast, reports on exacerbated airway neutrophilia and the implication of lung neutrophils as the main cell type driving ARDS in severe patients have yielded inconclusive results^7, 8, 17, 47^. Here, we present the first comprehensive study of the lung immune response to SARS-CoV-2 in Black/AA individuals and unequivocally identify a robust and sustained airway neutrophilia associated with disease severity. The COVID-19 pandemic further highlighted some of the socio-economic and behavioral inequalities that may have contributed to troubling disparities in COVID-19-associated morbidity and mortality^48^, with almost 70% of deaths being Black/AA patients in some areas^1, 2^. Although the socio-economic and behavioral differences indeed contribute to health disparities among demographics, a systematic investigation to determine the immunological features that characterize disease severity within Black/AA patients is lacking. Our systems biology approach addresses this knowledge gap and reveals new therapeutic targets to inhibit neutrophil migration, retention, and/or survival in the lung as potential effective interventions for individuals with severe disease that have been disproportionally affected by COVID-19.

Although previous studies on severe COVID-19 in humans have not been conclusive with regards to the role of airway neutrophilia in ARDS, studies in SARS-CoV-2-infected rhesus macaques and mice, where conditions and sample collection are more controlled, have identified exacerbated neutrophilia as a key immunological feature associated with disease severity^49, 50, 51^. Hence, taken together with previous studies on neutrophils^8, 14, 17, 45, 46, 47^ along with animal models^49, 50, 51^, our current findings provide compelling evidence that targeting exacerbated airway neutrophilia may constrain ARDS and prevent further lung damage in most patients requiring mechanical ventilator support. Although our studies represent one of the few targeted investigations in Black/AA patients, we believe our results have broader implications and may apply to most patients suffering from neutrophilic ARDS^11^.

The heightened numbers of neutrophils in the lung are likely to induce and sustain inflammatory signatures by an autocrine/paracrine feedback loop among neutrophils, and paracrine signaling to other cell types that can potentiate disease severity. Notably, NE is a potent serine protease that we show is released from degranulating neutrophils in the lungs (see Fig. 2), and has potential to stimulate production of TNF, IL-1β, and IL-8^52, 53^, and also abrogate protective effector functions of T cells and MdCs in the lung by cleaving cell surface receptors such as TLRs and Fc-receptors^54, 55^. Indeed, we demonstrate NE staining on the surface of T cells and MdCs in the lung (see Extended Data Fig. 5e) in addition to neutrophils, which may explain, in part, the reduction of FcγRII (CD32) expression on pulmonary neutrophils (see Extended Data Fig. 5a). Of note, the reduction in CD32 expression may prevent IgG-mediated suppression of ISG induction^14^ in pulmonary neutrophils. Indeed, we demonstrate a pronounced signature for the ISG *IFI30* (see Fig. 5f). Similarly, we show by intracellular staining that pulmonary neutrophils are producing exacerbated levels of IL-1β protein that is likely released upon degranulation along with IL-8 and IL-6 (see Figs. 2 and 3). In addition to neutrophils, the myeloid lineage is also abundant in the airways and as such they are poised to influence disease progression. For example, we and others^16^ have shown that infiltrating neutrophils sustain local production of calgranulins (S100A8/9/11/12), which can signal through and activate myeloid cells via TLRs and RAGE receptors, compounding the already hyperinflammatory lung milieu. Interestingly, the inflammatory monocytes that are likely recruited to the lungs via neutrophil-secreted CCL3/4 also show elevated levels of neutrophil chemotactic factor *CXCL8* (IL-8), which helps sustain recruitment of pathogenic neutrophils in a positive feedback loop.

Other immune and non-immune cell populations in the lung may contribute to COVID-19 pathogenesis in severe patients. Non-immune cells (e.g., stromal and epithelial cells) are known targets of SARS-CoV-2 in the lung^56^. Therefore, these cells are also posited to influence immune cell dynamics, especially at the outset of infection^57^. Indeed, our data indicate that non-immune cells play a role in granulocyte activation (Extended Data Fig. 5d) and therefore may initiate neutrophilia at the outset of infection^58^. Additionally, T cell-derived TNF and HMGB1, along with myeloid-derived IL-1β, have regulatory potential to induce inflammatory gene signatures observed in lung-recruited neutrophils. Strikingly, TNF is the ligand predicted to increase *BCL2A1* expression in pulmonary neutrophils, which is an anti-apoptotic factor known to regulate neutrophil survival^59, 60^. *TNF* signaling in recruited neutrophils also has the potential to drive *CCL4* (MIP-1β) expression, along with *CXCL16*, which is another chemotactic factor that can recruit neutrophils and macrophages via the CXCL16/CXCR6 pathway^61, 62^. In addition, myeloid-derived *APOE* is predicted to regulate the expression of *FCER1G* in lung-recruited neutrophils, which is associated with neutrophil activation during other viral respiratory infections^63^. Further studies should investigate other effector functions of myeloid and lymphoid subsets and how these cells interact with neutrophils to promote protection or pathology in the lungs of severe COVID-19 patients.

Interestingly, *CXCR4* is upregulated in recruited neutrophils in severe patients (Figs. 4d,i and 5f), which may promote neutrophil survival and retention during pneumonitis and/or further influence inflammatory neutrophil phenotype. Accordingly, CXCR4 was shown to promote transcriptional reprogramming of neutrophils in pulmonary tissues^64^. Although we do not find detectable levels of the canonical CXCR4 ligand (i.e., CXCL12) in the airways, we did observe transcripts for *HMGB1* (Fig. 6), which is an alternative ligand for CXCR4. Since the IL-8/CXCR2 pathway is most notably increased and likely the primary neutrophil recruitment axis to the airways, we speculate that CXCR4 signaling instead may promote neutrophil survival and retention at the site of inflammation, which has been previously reported^65^. Additionally, CXCR4 signaling can stimulate de novo *CXCL8*/IL-8 production^66, 67^, and has been shown to promote neutrophil extracellular trap release during malaria disease progression^68^. Alternatively, CXCR4 is also associated with neutrophil aging and senescence^69, 70, 71^. As such, elevated CXCR4 may be associated with prolonged neutrophil survival in COVID-19 pathogenesis. In a prior study^72^, we noted a similar pattern of CXCR4 expression on T cells from severe COVID-19 patients, where a progressive decrease of surface CXCR4 is associated with recovery. In contrast, patients that succumbed to the disease, showed a time-dependent escalation in CXCR4^+^ circulating T cells concomitant to increased CXCR4^+^ T cells in the lungs^72^, implicating CXCR4 on the dysregulated lung-homing inflammatory T cells in COVID-19. Further investigation into the potential role of CXCR4 in cell survival, retention, or recruitment may unravel novel therapeutic targets to modulate inflammation and treat severe COVID-19. Notably, therapeutic intervention with a CXCR4 antagonist during malaria-associated ARDS has shown significant benefit in animal models^36^.

Modulating inflammation through corticosteroids (particularly dexamethasone) has shown clinical efficacy and is now the standard-of-care for patients progressing to severe COVID-19^73^. However, the pleiotropic effects of glucocorticoids and their propensity to cause neutrophilia are well documented in asthma and chronic obstructive pulmonary disease (COPD)^74^. Although a well-established mechanism of action for dexamethasone is via transcriptional repression of proinflammatory cytokines, we observed very high *CXCL8* and *IL1B* transcripts concomitant with elevated pulmonary IL-8 and IL-1β protein with prominent signatures by intracellular staining—particularly in pulmonary neutrophils. In contrast, IL-6 levels were lower than that of IL-8 and IL-1β in our patient cohort, and overly abundant neutrophils were not producing as much IL-6 comparatively, which may explain, in part, why anti-IL-6/IL-6R studies failed to meet primary endpoints^75^. Here, we provide evidence of an uncoupled cytokine profile in airway fluids versus plasma where the lung microenvironment exhibits features of cytokine-induced ARDS driven largely by a proinflammatory, neutrophilic feed-forward loop. Beyond COVID-19, it has been shown that resolution of neutrophilia in ARDS has substantial prognostic benefit^10^. Taken together, the CXCL8 (IL-8)/CXCR2 signaling axis emerges as a key potential target for next-generation immunomodulatory therapy to reduce pathogenic neutrophilia and constrain severe disease in patients in addition to corticosteroids.

We also contend that recognizing the lung pathology in severe COVID-19 to be a neutrophilic and hyperinflammatory disease is paramount to achieve better outcomes in next-generation therapies. Although the lung pathology in COVID-19 is initiated by a viral infection, severe patients in the ICU no longer show signs of uncontrolled viral replication. In fact, not only did we not detect viral transcripts by scRNA-seq within the cells in the airways, but we also noted decreased viral burden in severe patients in the ICU versus mild-acute patients seen in the outpatient clinic (Fig. 7). This is further supported by our previous study where we performed plaque assays on the respiratory secretions from severe patients and revealed significantly diminished, if any, viral plaques from the endotracheal aspirates^76^. This may explain, in part, why antiviral drugs such as remdesivir are not able to prevent death when administered to severe patients in ICU^77^.

**Figure 7.**
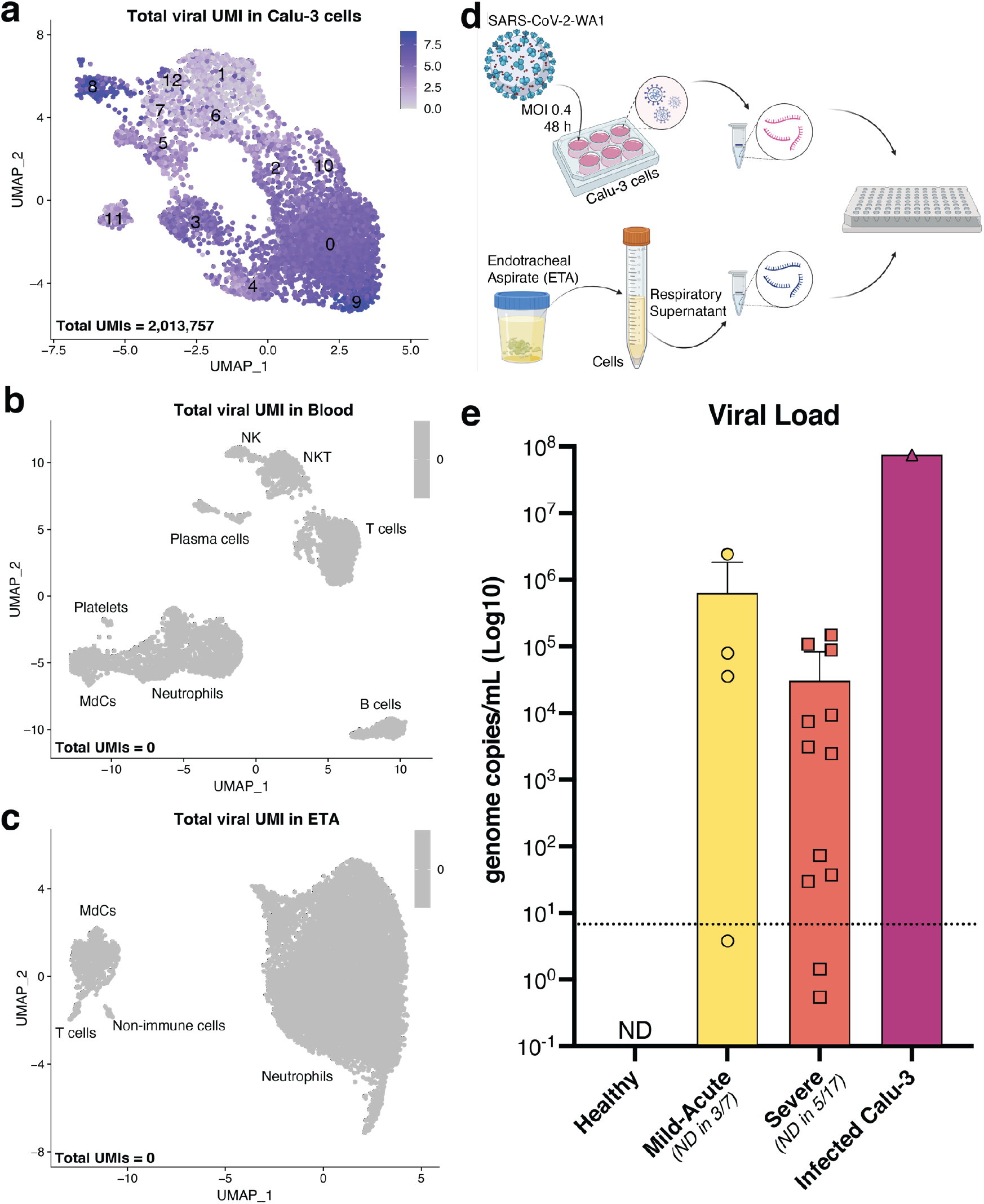
Viral transcripts (scRNA-seq) and viral load in the airways of severe patients. Calu-3 cells infected with SARS-CoV-2 USA-WA1/2020 (MOI 0.4) were encapsulated for scRNA-seq assays (a) following the same protocol used for patient cells from blood (b) and ETA (c). Millions of viral transcripts (or unique molecular identifiers; UMIs) were detected in about 5,000 Calu-3 cells (a), but not in the cells from severe patients (total UMI = 0; b and c). RT-qPCR (d) results of culture supernatant from Calu-3 infected cells and respiratory supernatants from mild-acute and severe patients (e) show a lower viral load in severe patients than in mild-acute patients and in the Calu-3-infected culture supernatant. ND = not detected; horizontal dotted line = lower limit of detection (LLOD).

It is important to consider potentially confounding factors and limitations in our study. Though a major strength of our study is the uniformity of sample collection in a single disease state, our airway samples are limited to ETA, which contrasts with bronchoalveolar lavage fluid (BALF) used in other studies^7, 17^. The ETA procedure can sample material from the medial airways— i.e., an intermediate between the distal/lower airways (e.g., BALF) and the proximal/upper airways (e.g., sputum or oro-/naso-pharyngeal samples). Although the relative abundance of immune cells may vary across the upper and lower airways, previous studies observe a correlation between paired ETA and BALF samples. In addition, these studies show that ETA samples are not inherently neutrophilic^78^, though neutrophilia/granulocytosis may be a shared feature of non-COVID-19 and COVID-19 ARDS^10, 79^. In any event, our study is one of the only known to date to employ integrated multi-omics single-cell investigation of immunity in exclusively Black/AA subjects, which are disproportionately burdened with severe disease and worse outcomes. Importantly, other studies of COVID-19 immune responses have noted similar features of disease severity^7, 15, 16^, which suggests that the findings in our target demographic will be broadly applicable to other groups. Finally, our study was limited in scope to 35 total subjects and single collection time points, limiting our abilities to interrogate correlations with clinical outcomes, warranting further longitudinal studies in larger patient cohorts.

In conclusion, we present evidence that neutrophils are poised to be the leukocyte population most responsible for the dysregulated hyperinflammatory response that drives ARDS in severe COVID-19 patients. Neutrophil frequency and inflammatory profiles reveal that they are not only the most abundant leukocyte population in the medial airways, but also major producers of hallmark effector molecules associated with disease severity, including IL-8, IL-1β, and IL-6, along with potent proteases such as NE and MPO, which are inherently inflammatory and contribute to lung damage/pathology. Furthermore, we provide evidence for a neutrophil feedforward loop where IL-8, produced by virtually all pulmonary neutrophils (and some myeloid and stromal cells), is the primary chemokine recruiting circulating neutrophils and promoting neutrophilia in the inflamed airways.

Collectively, our findings implicate neutrophilia in the immunopathophysiology of severe COVID-19 disease where perpetual, transcriptionally active, and highly inflammatory pulmonary neutrophils drive ARDS despite low viral burden. Thus, therapeutic intervention targeting neutrophil recruitment/retention and/or survival/reprogramming at the site of inflammation has the potential to constrain ARDS in severe patients, particularly those most vulnerable to succumb to COVID-19 disease.

## Methods

### Ethics and biosafety

A total of 35 individuals were recruited for this study (Extended Data Tables 1 and 2). The 18 severe COVID-19+ patients were recruited from the Intensive Care Units of Emory University, Emory St. Joseph’s, Emory Decatur, and Emory Midtown Hospitals. We also recruited 9 mild COVID-19-infected outpatients in the Emory Acute Respiratory Clinic and 8 healthy adults from the Emory University Hospital. All studies were approved by the Emory Institutional Review Board (IRB) under protocol numbers IRB00058507, IRB00057983, and IRB00058271. Informed consent was obtained from the patients when they had decision-making ability or from a legally authorized representative (LAR) if the patient was unable to provide consent. Blood and sputum or endotracheal aspirate (ETA) were obtained. Control blood samples were obtained from healthy adults matched by age and race. Study inclusion criteria included a confirmed COVID-19 diagnosis by PCR amplification of SARS-CoV-2 viral RNA obtained from nasopharyngeal or oropharyngeal swabs, age of 18 years or greater, and willingness to provide informed consent. Individuals with a confirmed history of COVID-19 diagnosis were excluded from the healthy donor group. All work with infectious virus and respiratory samples from COVID-19 patients was conducted inside a biosafety cabinet within the Emory Health and Safety Office (EHSO) and the United States Department of Agriculture (USDA)-approved BSL3 containment facility in the Health Sciences Research Building at Emory University following protocols approved by the Institutional Biosafety Committee (IBC) and Biosafety Officer (see ref^76^).

### Patient sample collection and processing

Primary leukocytes from the airways of COVID-19 patients requiring mechanical ventilator support were collected bedside via endotracheal aspiration (ETA) and whole blood collected by standard venipuncture. Plasma from whole blood was isolated via centrifugation at 400 × g for 10 min at 4°C. To remove platelets, the isolated plasma was centrifuged at 4,000 × g for 10 min at 4°C. Untouched circulating leukocytes were isolated using the EasySep™ RBC Depletion Reagent (StemCell Technologies). ETA (from severe patients) or non-induced sputum (from mild patients) was mixed 1:1 with a 50mM EDTA solution (final concentration 25 mM EDTA) in custom RPMI-1640 media deficient in biotin, L-glutamine, phenol red, riboflavin, and sodium bicarbonate (defRPMI-1640), with 3% newborn calf serum (NBCS) and mechanically dissociated using a syringe to liberate leukocytes from mucins and other respiratory secretions. Supernatants were collected for further analysis, and then cells underwent an additional mechanical dissociation step using 1-3 mL of a 10 mM EDTA in defRPMI-1640 + 3% NBCS and a P1000 pipettor. Cells were then washed with 10 mL defRPMI-1640 + 3% NBCS, passed through a 70 µm nylon strainer, and pelleted through a 2 mL 100% NBCS layer prior to counting and downstream processing.

### High-dimensional (Hi-D) 30-parameter flow cytometry

Cells (up to 10^7^ total) were resuspended in defRPMI-1640 with 3% newborn calf serum and Benzonase™ (FACS buffer) in 5 mL FACS tubes and pre-incubated with GolgiStop™ (BD Biosciences) for ∼60 min at 4°C. Human TruStain FcX™ was then added, followed by a 10 min incubation at RT. The 24-color extracellular staining master mix (Extended Data Table 2) was prepared 2X in BD Horizon™ Brilliant Stain Buffer to prevent staining artifacts from BD Horizon Brilliant dye interactions and added 1:1 to cells, then incubated for 30 min at 4°C. Following staining (and total 1 h exposure to GolgiStop™), cells were washed with ∼4 mL FACS buffer. Next, the cells were resuspended in 200 µL of BD Cytofix/Cytoperm™ fixation/permeabilization solution and incubated at 4°C for 30 min followed by a wash with ∼4 mL BD Perm/Wash™ Buffer. The 4-color intracellular staining (Extended Data Table 3) was prepared in BD Perm/Wash™ Buffer, and cells were stained for 30 min at 4°C. Cells were washed with ∼4 mL BD Perm/Wash™ Buffer, then resuspended for a final 20 min incubation in 4% PFA and transported out of the BSL3 containment facility. Cells were washed in ∼4 mL FACS buffer, then resuspended in 200-1000 µL FACS buffer for acquisition using BD FACSDiva™ Software on the Emory Pediatric/Winship Flow Cytometry Core BD FACSymphony™ A5. To distinguish auto-fluorescent cells from cells expressing low levels of a particular surface marker, we established upper thresholds for auto-fluorescence by staining samples with fluorescence-minus-one (FMO) control stain sets in which a reagent for a channel of interest is omitted. Data were analyzed with FlowJo™ v10.8 (FlowJo LLC).

### Cell-surface antibody-derived tag (ADT) staining, single-cell encapsulation, and library generation

Leukocytes from whole blood and ETA samples were incubated with oligo-conjugated Ig-A/D/G/M for 10 min at 4°C followed by the addition of Human TruStain FcX™ (BioLegend) and 10 min incubation at RT. Cells were then surface stained with oligo-conjugated monoclonal antibody panel (total 89 antibodies; Extended Data Table 5) for 30 min at 4°C, followed by two washes in def-RPMI-1640/0.04% BSA. Cells were resuspended at a concentration of 1200-1500 cells/µL in def-RPMI-1640/0.04% BSA and passed through a 20 or 40 µm cell strainer before loading onto a Chromium Controller (10X Genomics, Pleasanton, CA). Cells were loaded to target encapsulation of 10,000 cells. Gene expression (GEX) and antibody-derived tag (ADT) libraries were generated using the Chromium Single Cell 5′ Library & Gel Bead Kit v1.1 with feature barcoding following the manufacturer’s instructions. GEX libraries were pooled and sequenced at a depth of approximately 540,000,000 reads per sample in a single S4 flow cell and ADT libraries at a depth of approximately 79,000,000 reads per sample in a single lane of an S4 flow cell on a NovaSeq™ 6000 (Illumina, San Diego, CA; Extended Data Table 6)

### Multi-omics single-cell RNA sequencing (scRNA-seq) analysis

Single-cell 5′ unique molecular identifier (UMI) counting and barcode de-multiplexing were performed using CellRanger Software (v.5.0.0). To detect SARS-CoV-2 viral RNA reads, we built a custom reference genome from human GRCh38 and SARS-CoV-2 references (severe acute respiratory syndrome coronavirus 2 isolate Wuhan-Hu-1, complete genome, GenBank MN908947.3). Splicing-aware aligner STAR^80^ was implemented to align FASTQ inputs to the reference genome, and the resulting files are automatically filtered by CellRanger to include only cell barcodes representing real cells. This determination is based on the distribution of UMI counts. ADT reads were aligned to a feature reference file containing the antibody-specific barcode sequences. To recover neutrophils, we applied our SuPERR-seq pipeline as previously described^30^. Briefly, we recovered neutrophils from CellRanger unfiltered count matrices by plotting surface CD16 ADT and CD66b ADT using the “FeatureScatter” function in Seurat v4.0^81^ (R version 4.0.2). The double-positive cell barcodes were then extracted and further evaluated by GEX to confirm viable neutrophil identity. A threshold for mitochondrial content per barcode was determined for each sample independently and applied as a cutoff to remove dead or dying cells (Extended Data Table 7). Most samples show high cell viability with a minimal proportion of dead cells.

The UMI counts of the GEX data were log-normalized by the “NormalizeData” function in Seurat before downstream analysis, following the optimal workflow we previously described for sample normalization and data integration^82^. Center log-ratio (CLR) transform in Seurat was performed on ADT UMIs when recovering neutrophils from the unfiltered matrices. For surface protein visualization to classify major lineages using our SuPERR-seq workflow^30^, ADT UMIs were normalized using the R package Denoised and Scaled by Background^83^ (DSB) to remove ambient UMI counts (i.e., background) prior to manual sequential gating by surface expression (Extended Data Fig. 4) in SeqGeq v1.7 (FlowJo, LLC). DSB uses empty droplets to calculate background expression, which was manually selected according to the distribution of total ADT per cell in the raw count matrices (Extended Data Table 8). To minimize the influence from non-informative empty droplets, we removed cell barcodes with less than 100 total ADT UMIs before plotting the ADT distribution.

Before integrating the multiple datasets, we first classified major lineages in individual samples based on a combination of gene transcript and surface protein markers (SuPERR-seq workflow^30^) as in Fig. 4 for samples where the ADT library was of sufficient quality to allow manual gating (Extended Data Fig. 4). Cell barcodes within each major lineage that co-expressed markers exclusive to other major lineages were considered cell doublets and removed (Extended Data Fig. 4). In addition, we removed cell barcodes with extremely high total ADT UMIs, which we considered to be aggregated cells. To efficiently integrate replicate samples, we concatenated major lineages derived from the same tissue in different donors. To minimize batch effects and optimize data integration, we followed the data normalization and merging strategies described previously^82^. Briefly, samples were first treated individually, and log-normalized count matrices were scaled/Z-transformed, and the “vst” method of the Seurat function “FindVariableFeatures” was utilized to select the top 1000 highly variable genes (HVGs) of each sample. HVGs shared between replicate samples were used to perform principal component analysis (PCA). To visualize the data, we performed UMAP reduction of the first 30 PCs, and cell clustering was generated using the Leiden community detection algorithm at a resolution of 0.8. UMAP visualizations for the integrated Blood and integrated ETA were generated using Seurat v4 data integration workflow.

### Receptor-ligand interaction analyses

Clustered cells from lung and blood samples from each patient were investigated for evidence of intercellular communication using CellChat^31^. Clustered cell populations from the lung samples were combined with the blood neutrophils to determine which lung cell populations could recruit circulating neutrophils. We utilized NicheNet^38^ to determine which ligand-receptor pairs could be responsible for the different transcriptional states of the neutrophil populations in the lung and blood samples. We focused on certain neutrophil clusters as the receiver populations, considering the remaining neutrophils and other lung cell populations as the senders and thus potential interactors. The target set of genes was determined using Seurat::FindMarkers(min.pct = 0.1), keeping only those genes with an adjusted p-value lower than 0.05 and average log2-fold change of more than 0.2. To address gene expression changes due to lung infiltration, we ran the algorithm using blood neutrophil cluster 2 as the most likely candidate for lung infiltration. The genes considered were those differentially expressed between blood cluster 2 and the lung neutrophils and genes differentially expressed between blood cluster 2 and those cells that progress along “Trajectory 2” in the lung (see Fig.5).

### Cell trajectory analyses

The Python toolkit scVelo^32^ inferred the trajectories using biological data of the ETA neutrophils. Input data for scVelo analysis was intron, exon, and spanning count matrices estimated using the dropEST tool^84^, then filtered with previously identified neutrophil cell barcodes in R studio. Intron, exon, and spanning matrices were compared to identify missing rows (genes) and were added to each matrix to equalize dimensions. The exon matrix contained the spliced matrix, and the sum of the intron and spanning matrices constituted the unspliced matrix. Spliced and unspliced matrices were imported with anndata library, and pandas library was used to import gene names and cell barcodes. Raw count matrices were added to anndata object layers as ‘spliced’ and ‘unspliced.’ Then, gene names and cell barcodes were attached to variables and observations of anndata object, respectively. Anndata object was transposed, followed by the regular scVelo analysis. The default parameters of plotting velocity streams include vkey=‘velocity’, colorbar=True, alpha=0.3, sort_order=True, and legend_loc=‘on data’.

### Pathway and process enrichment analyses

Differential gene expression (DGE) analyses were performed in Seurat v4 and imported for gene annotation and further analysis using Metascape^85^. DGE between neutrophils versus the total ETA were used to generate Fig. 5g and non-immune cells versus total ETA were used for Extended Data Fig. 5d.

### Mesoscale U-PLEX assays

U-PLEX Biomarker Group 1 Human Multiplex Assays (Meso Scale Discovery) were used to evaluate levels of 21 analytes following the manufacturer’s protocol (Extended Data Table 4) in plasma and UVC-inactivated respiratory supernatants (see ref^76^). Samples were diluted 1:5 for all assays except for IL-8, MCP-1, and IL-1RA, which were above the upper limit of detection for the assay and were diluted 1:200 to acquire measurement within the assay range. Final values were obtained by multiplying measurements by their respective dilution factor.

### Myeloperoxidase (MPO) content and activity

The abundance and activity of MPO were quantified as previously described^29^. MPO activity and protein concentration were measured sequentially following the immunocapture. On average, across six 96-well plate assays, lower limits of quantification were 4.0 ng/mL (activity) and 0.84 ng/mL (protein). Samples above the lower limit of detection but below the lower limit of quantification were imputed as half of the latter, and those detected above the highest standard of 50 ng/mL (i.e., above the upper limit of detection) at all dilutions were imputed as twice the standard concentration (Extended Data Table 4).

### SARS-CoV-2 quantitative reverse transcription PCR (RT-qPCR)

Viral (v)RNA was extracted from the respiratory secretions of COVID-19 patients using the *Quick*-RNA™ Viral Kit (Zymo Research) following the manufacturer’s protocol and complementary (c)DNA synthesized using the High-Capacity cDNA Reverse Transcription Kit (Applied Biosystems™) per the manufacturer’s instructions, then diluted 1:5 in nuclease-free water. 10 µL diluted cDNA was used with the NEB Luna^®^ Universal Probe qPCR Master Mix (New England BioLabs^®^ Inc.) following the manufacturer’s protocol and performed in 384-well plates using a QuantStudio™ 5 Real-Time PCR System (Applied Biosystems™). Primer/probe pairs were: AGAAGATTGGTTAGATGATGATAGT (forward), TTCCATCTCTAATTGAGGTTGAACC (reverse), and /56-FAM/TCCTCACTGCCGTCTTGTTGACCA/3IABkFQ/ (probe), which were designed from sequences previously described^86^ (Integrated DNA Technologies; IDT). To generate a standard curve for the quantification of SARS-CoV-2 genome copies a gBlock from IDT with the following sequence was used as a standard: AATTAAGAACACGTCACCGCAAGAAGAAGATTGGTTAGATGATGATAGTCAACAAACTGTTGGTCAACAAGACGGCAGTGAGGACAATCAGACAACTACTATTCAAACAATTGTTGAGGTTCAACCTCAATTAGAGATGGAACTTACAGTTTCAGTGTTCAATTAA.

### Statistical analyses

Statistical analyses were performed using GraphPad Prism9. Data were analyzed for distribution (normal (Gaussian) vs. lognormal) independently using the D’Agostino and Pearson test for normality in the untransformed and Log10-transformed data. In cases where the sample size (*N*) was too small for D’Agostino and Pearson normality test, the Shapiro-Wilk test was used to assess distribution. When data passed both distribution tests, the likelihood of each distribution (normal vs. lognormal) was computed, and QQ-plots were generated. When Log10 transformed data had a higher likelihood of a normal distribution (passing normal distribution test) and/or failed lognormal distribution test, paired t-tests were performed to compare matching blood and respiratory supernatant samples within a single group. If the data had unequal variance (as determined by an F-test), a ratio paired t-test was performed. All instances where lognormal distribution was likely non-parametric Wilcoxon matched-pairs sign ranked tests were performed. For comparisons across the three patient groups (i.e., healthy, mild-acute, severe), ordinary one-way ANOVA (if equal variance) or Brown-Forsythe and Welch ANOVA (if unequal variance) tests were performed for data with a normal distribution. Alternatively, data with a lognormal distribution were analyzed with a Kruskal-Wallis test.

## Data availability

Single-cell sequencing datasets presented here are available through NCBI GEO, accession number XXX.

## Author Contributions

**Devon J. Eddins**: Conceptualization, Methodology, Investigation, Formal Analysis, Data Curation, Writing–Original Draft, Writing–Review & Editing, Visualization. **Junkai Yang**: Methodology, Formal Analysis, Data Curation, Writing–Review & Editing, Visualization. **Astrid Kosters**: Methodology, Investigation, Data Curation, Writing–Review & Editing. **Vincent D. Giacalone**: Methodology, Formal Analysis, Investigation, Data Curation, Writing–Review & Editing. **Ximo Pechuan**: Methodology, Formal Analysis, Data Curation, Visualization, Writing– Review & Editing. **Joshua D. Chandler**: Methodology, Formal Analysis, Investigation, Data Curation, Writing–Review & Editing. **Jinyoung Eum**: Formal Analysis, Visualization. **Benjamin R. Babcock**: Investigation, Data curation. **Brian S. Dobosh**: Investigation, Data Curation, Formal Analysis. **Mindy R. Hernández**: Resources. **Fathma Abdulkhader**: Investigation. **Genoah L. Collins**: Investigation. **Richard P. Ramonell**: Resources, Data Curation, Writing– Review & Editing. **Christine Moussion**: Methodology, Formal Analysis. **Darya Y. Orlova**: Methodology, Data Curation. **Ignacio Sanz**: Resources. **F. Eun-Hyung Lee**: Methodology, Data Curation, Resources, Writing–Review & Editing. **Rabindra M. Tirouvanziam**: Methodology, Formal Analysis, Data Curation, Resources, Writing–Review & Editing. **Eliver E**.**B. Ghosn**: Conceptualization, Methodology, Formal Analysis, Resources, Data Curation, Writing–Original Draft, Visualization, Supervision, Project Administration, Funding Acquisition. All authors discussed the results and read and approved the final manuscript.

## Acknowledgments

This study was funded by NIH/NIAID R01AI123126 (EEBG) and R01AI123126-05S1 (EEBG), the Program for Breakthrough Biomedical Research (EEBG), and the Lowance Center for Human Immunology (EEBG). DJE was partially supported by the Emory’s Laney Graduate School Fellowship, and RPR was supported by the NIH T32-HL116271-07 Fellowship. The graphical abstract and Fig. 7d were generated in part using BioRender. We thank Keivan Zandi, Ann Chahroudi, Nils Schoof, Kira Moresco, and Stacy Heilman of the Department of Pediatrics (Emory University) along with the Emory Biosafety Officers Kalpana Rengarajan and Esmeralda Meyer for their assistance with setting up the BLS3 facility and kindly providing SARS-CoV-2 viral stocks. We thank Nadia Roan (Gladstone Institutes/UCSF) and Sulggi Lee (UCSF) for helpful discussions. Flow cytometry data were collected at the Emory’s Pediatrics/Winship Flow Cytometry Core (access supported in part by Children’s Healthcare of Atlanta). We acknowledge the Genomic Cores at Yerkes Non-Human Primate Research Center at Emory University (NIH P51 OD011132; NIH S10 OD026799), Baylor College of Medicine, and the Parker H. Petit Institute for Bioengineering and Bioscience at the Georgia Institute of Technology for the single-cell library sequences. We are grateful for the efforts of Sang N. Le, John Varghese, Anum Jalal, Saeyun Lee, and Rahul Patel, who also contributed to patient recruitment. We also thank the nurses, staff, and providers in the 71 ICU in Emory University Hospital Midtown, the medical ICU in Emory Decatur Hospital, the 5G/6G ICU in Emory University Hospital, and the ICU in Emory Saint Joseph’s Hospital for their dedication and commitment during the COVID-19 pandemic. We thank all healthy individuals, patients, and their families for their participation in this study, without whom our work would not have been possible.

## Supplemental Material

**Extended Data Table 1.** Demographic and clinical data from the 35 Black/African American individuals enrolled in our studies. Table describing demography, status, sex, and age of study participants along with select details of clinical course including administration of Dexamethasone and Remdesivir, date first intubated (severe, ICU patients only), date of PCR positive diagnosis (Dx), date clinical specimens were collected, and total length ICU stay in days, and clinical outcome (whether recovered or succumbed to disease).

| # | Demographic | PID | Status | Sex | Age Group | Dexamethasone or equivalent | Remdesivir (No. of doses) | Date PCR positive Dx | Date intubated | Date sample collected | Length ICU stay (days) | Clinical Outcome |
| --- | --- | --- | --- | --- | --- | --- | --- | --- | --- | --- | --- | --- |
| 1 | AA | PHA0001 | Severe | Male | 35-54 | Yes | 0 | 6/9/2020 | 6/8/2020 | 6/17/2020 | 32 | Recovered |
| 2 | AA | PHA0004 | Severe | Male | 35-54 | Yes | 0 | 7/11/2020 | 7/18/2020 | 8/5/2020 | 37 | Recovered |
| 3 | AA | PHA0006 | Severe | Male | 35-54 | Yes | 0 | 8/11/2020 | 8/16/2020 | 8/20/2020 | 8 | Recovered |
| 4 | AA | PHA0018 | Severe | Male | 35-54 | Yes | 2 | 10/14/2020 | 9/29/2020 | 10/8/2020 | 13 | Recovered |
| 5 | AA | PHA0041 | Severe | Male | 35-54 | Yes | 2 | 2/6/2021 | 2/6/2021 | 2/10/2021 | 14 | Recovered |
| 6 | AA | PHA0003 | Severe | Male | 55+ | Yes | 2 | 6/23/2020 | 6/25/2020 | 6/30/2020 | 14 | Succumbed |
| 7 | A | PHA0008 | Severe | Male | 55+ | Yes | 2 | 8/29/2020 | 8/30/2020 | 9/3/2020 | 19 | Recovered |
| 8 | AA | PHA0024 | Severe | Male | 55+ | Yes | 2 | 12/27/2020 | 12/31/2020 | 1/6/2021 | 9 | Recovered |
| 9 | AA | PHA0025 | Severe | Male | 55+ | Yes | 0 | 12/20/2020 | 12/24/2020 | 1/8/2021 | 26 | Recovered |
| 10 | AA | PHA0031 | Severe | Male | 55+ | Yes | 1 | 1/23/2021 | 1/24/2021 | 1/27/2021 | 11 | Succumbed |
| 11 | AA | PHA0032 | Severe | Male | 55+ | Yes | 1 | 1/9/2021 | 1/14/2020 | 1/27/2021 | 40 | Recovered |
| 12 | AA | PHA0035 | Severe | Male | 55+ | Yes | 0 | 1/18/2021 | 1/20/2021 | 2/2/2021 | 53 | Recovered |
| 13 | AA | PHA0036 | Severe | Female | 18-34 | Yes | 2 | 1/10/2021 | 12/31/2020 | 2/2/2021 | 42 | Recovered |
| 14 | AA | PHA0002 | Severe | Female | 55+ | Yes | 0 | 6/20/2020 | 6/20/2020 | 6/24/2020 | 25 | Recovered |
| 15 | AA | PHA0005 | Severe | Female | 55+ | Yes | 0 | 7/16/2020 | 7/18/2020 | 8/12/2020 | 33 | Recovered |
| 16 | AA | PHA0007 | Severe | Female | 55+ | Yes | 0 | 8/9/2020 | 8/10/2020 | 9/2/2020 | 60 | Recovered |
| 17 | AA | PHA0010 | Severe | Female | 55+ | Yes | 2 | 9/17/2020 | 9/21/2020 | 9/24/2020 | 18 | Recovered |
| 18 | AA | PHA0026 | Severe | Female | 55+ | Yes | 1 | 12/27/2020 | 12/28/2020 | 1/8/2021 | 16 | Succumbed |
| 19 | A | PHA0040 | Mild-Acute | Male | 35-54 | Yes | 0 | 1/6/2021 | N/A | 2/8/2021 | N/A | Recovered |
| 20 | AA | PHA0043 | Mild-Acute | Male | 35-54 | Yes | 0 | Unclear | N/A | 2/18/2021 | N/A | Recovered |
| 21 | AA | PHA0023 | Mild-Acute | Male | 55+ | Yes | 0 | 11/6/2020 | N/A | 11/11/2020 | N/A | Recovered |
| 22 | AA | PHA0047 | Mild-Acute | Male | 55+ | Yes | 0 | 2/8/2021 | N/A | 2/19/2021 | N/A | Recovered |
| 23 | AA | PHA0012 | Mild-Acute | Female | 35-54 | Yes | 0 | 9/18/2020 | N/A | 9/29/2020 | N/A | Recovered |
| 24 | AA | PHA0013 | Mild-Acute | Female | 35-54 | Yes | 0 | 9/21/2020 | N/A | 9/29/2020 | N/A | Recovered |
| 25 | AA | PHA0044 | Mild-Acute | Female | 35-54 | Yes | 0 | 2/16/2021 | N/A | 2/19/2021 | N/A | Recovered |
| 26 | AA | PHA0046 | Mild-Acute | Female | 35-54 | Yes | 0 | 2/12/2021 | N/A | 2/19/2021 | N/A | Recovered |
| 27 | AA | PHA0045 | Mild-Acute | Female | 55+ | Yes | 0 | 2/12/2021 | N/A | 2/19/2021 | N/A | Recovered |
| 28 | AA | PHA1009 | Healthy | Female | 18-34 | Yes | 0 | N/A | N/A | 10/23/2020 | N/A | N/A |
| 29 | AA | PHA1005 | Healthy | Female | 35-54 | Yes | 0 | N/A | N/A | 10/23/2020 | N/A | N/A |
| 30 | AA | PHA1006 | Healthy | Female | 35-54 | Yes | 0 | N/A | N/A | 10/23/2020 | N/A | N/A |
| 31 | AA | PHA1007 | Healthy | Female | 35-54 | Yes | 0 | N/A | N/A | 10/23/2020 | N/A | N/A |
| 32 | AA | PHA1008 | Healthy | Female | 35-54 | Yes | 0 | N/A | N/A | 10/23/2020 | N/A | N/A |
| 33 | AA | PHA1010 | Healthy | Female | 35-54 | Yes | 0 | N/A | N/A | 10/23/2020 | N/A | N/A |
| 34 | AA | PHA1011 | Healthy | Female | 35-54 | Yes | 0 | N/A | N/A | 2/10/2021 | N/A | N/A |
| 35 | AA | PHA1012 | Healthy | Female | 55+ | Yes | 0 | N/A | N/A | 2/17/2021 | N/A | N/A |

**Extended Data Table 2.** Clinical characteristics of the 35 Black/African American individuals enrolled in our studies. Table describing clinical characteristics of 35 patients across the 3 cohorts (healthy, mild-acute, and severe) including age, gender, site of endotracheal aspiration (ETA), medical history, along with complications for the severe patients admitted to the ICU.

| Characteristics | All subjects (N = 35) |  |  |
| --- | --- | --- | --- |
|  | Healthy<br>(N = 8) | Mild-Acute<br>(N=9) | Severe<br>(N=18) |
| **Age** |  |  |  |
| A. Mean age ± SD | 46 ± 11 | 51 ± 12 | 56 ± 14 |
| B. Age range | 31 – 70 | 35 – 68 | 22 – 81 |
| **Gender** |  |  |  |
| A. Female | 8 (100%) | 5 (56%) | 6 (33%) |
| B. Male | 0 (0%) | 4 (44%) | 12 (67%) |
| **Site of endotracheal aspiration** |  |  |  |
| A. Endotracheal tube | — | — | 9 (50%) |
| B. Tracheostomy | — | — | 9 (50%) |
| **Medical History** |  |  |  |
| A. Smoking History |  |  |  |
| i) Never | 7 (87.5%) | 7 (78%) | 12 (67%) |
| ii) Former | 1 (12.5%) | 2 (22%) | 2 (11%) |
| iii) Current | 0 (0%) | 0 (0%) | 4 (22%) |
| B. Hypertension (HTN) | 3 (37.5%) | 3 (33%) | 11 (61%) |
| C. Obesity | 2 (25%) | 1 (11%) | 4 (22%) |
| D. Asthma | 2 (25%) | 2 (22%) | 2 (11%) |
| E. COPD | 0 (0%) | 0 (0%) | 2 (11%) |
| F. Diabetes mellitus (DM) |  |  |  |
| i) Type I DM | 0 (0%) | 0 (0%) | 0 (0%) |
| ii) Type II DM | 1 (12.5%) | 0 (0%) | 8 (44%) |
| G. Deep vein thrombosis (DVT) | 0 (0%) | 0 (0%) | 1 (5%) |
| H. Hypothyroidism | 0 (0%) | 0 (0%) | 1 (5%) |
| I. Congestive Heart Failure (CHF) | 1 (12.5%) | 0 (0%) | 5 (28%) |
| J. Gastroesophageal reflux disease (GERD) | 1 (12.5%) | 1 (11%) | 1 (5%) |
| K. Dyslipidemia | 0 (0%) | 1 (11%) | 1 (5%) |
| L. Obstructive sleep apnea (OSA) | 0 (0%) | 0 (0%) | 2 (11%) |
| M. Coronary artery disease (CAD) | 0 (0%) | 0 (0%) | 1 (5%) |
| N. Chronic kidney disease | 0 (0%) | 1 (11%) | 1 (5%) |
| O. Cancer | 0 (0%) | 0 (0%) | 0 (0%) |
| P. Autoimmune hemolytic anemia | 0 (0%) | 0 (0%) | 1 (5%) |
| Q. Thrombocytopenia | 0 (0%) | 0 (0%) | 1 (5%) |
| R. Arthritis/Rheumatoid arthritis (RA) | 1 (12.5%) | 0 (0%) | 1 (5%) |
| **Complications** |  |  |  |
| A. Acute respiratory distress syndrome (ARDS) | — | — | 18 (100%) |
| B. Hypoxemic respiratory failure | — | — | 13 (72%) |
| C. Extracorporeal membrane oxygenation (ECMO) | — | — | 2 (11%) |
| D. Shock | — | — | 15 (83%) |
| E. Nosocomial pneumonia | — | — | 6 (33%) |
| F. Coagulopathy | — | — | 3 (17%) |
| G. Seizures | — | — | 3 (17%) |
| H. Cardiac arrhythmia | — | — | 3 (17%) |
| I. Renal failure | — | — | 8 (44%) |
| J. Renal replacement therapy | — | — | 6 (33%) |

**Extended Data Table 3.**
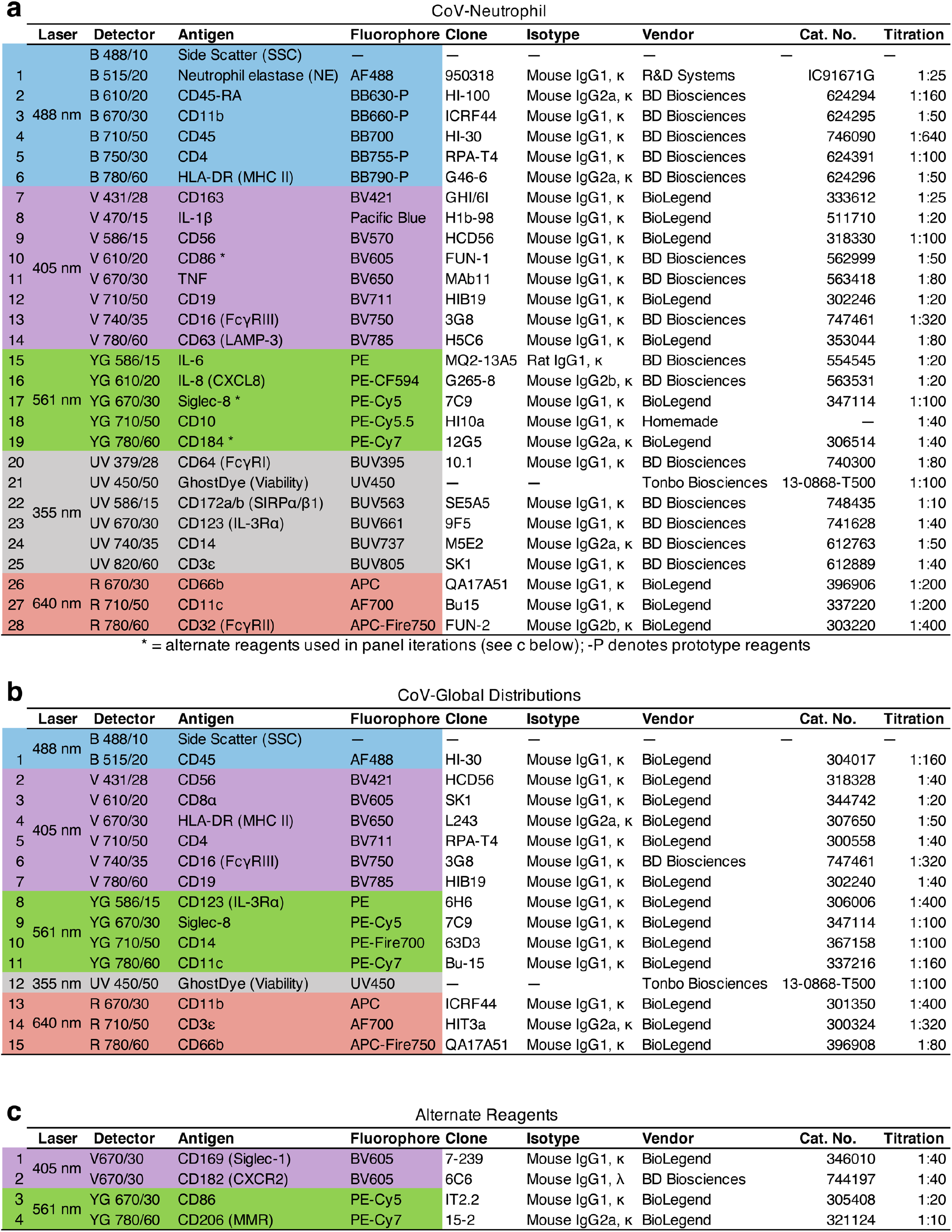
High-dimensional 30-parameter, including intracellular cytokine staining, and 17-parameter flow cytometry panels used for airway and blood cells. Flow cytometer configuration and cytometry reagents used in the final panels to interrogate (a) neutrophil phenotype and (b) global immune cell distributions and blood and ETA samples. Channels marked with an asterisk (*) denote alternate reagents (c) used in earlier panel iterations for some samples in this study. Monoclonal antibody (mAb) master mixes were prepared in BD Horizon™ Brilliant Stain Buffer and samples stained as described in the methods section. Titrations of all reagents were determined in house for each lot independently prior to use. AF: AlexaFluor, APC: Allophycocyanin, BB: Brilliant Blue, BUV: Brilliant Ultraviolet, BV: Brilliant Violet, FITC: Fluorescein isothiocyanate, PE: Phycoerythrin. Fluorophores marked with -P denote prototype reagents and are custom conjugations from BD Biosciences.

**Extended Data Table 4.** Mesoscale U-PLEX biomarker group 1 human and myeloperoxidase (MPO) assays. Median ± SD and range of lower limits of detection (LLOD) for (a) Mesoscale UPLEX analytes and (b) MPO assays. Lower limits of quantification (LLOQ; MPO assays) and LODs (determined by standard curve) were run for each assay plate independently and the median LLOD plotted as horizontal dotted lines for each assay (see Fig. 3 and Extended Data Fig. 2).

| **a** | **Analyte** | **Median LLOD**<br>**(pg/mL)** | **LLOD Range**<br>**(pg/mL)** |
| --- | --- | --- | --- |
| 1 | G-CSF (CSF3) | 0.84 ± 0.88 | 0.15—2.12 |
| 2 | GM-CSF (CSF2) | 0.11 ± 0.13 | 0.01—0.31 |
| 3 | IFN-γ | 1.31 ± 1.12 | 0.16—2.79 |
| 4 | IL-10 | 0.03 ± 0.02 | 0.01—0.06 |
| 5 | IL-1α | 0.13 ± 0.07 | 0.04—0.19 |
| 6 | IL-1β | 0.07 ± 0.05 | 0.01—0.12 |
| 7 | IL-6 | 0.31 ± 0.39 | 0.03—0.89 |
| 8 | IL-8 (CXCL8) | 0.03 ± 0.03 | 0.02—0.08 |
| 9 | TNF | 0.24 ± 0.13 | 0.07—0.35 |
| 10 | VEGFA | 0.43 ± 0.44 | 0.06—1.06 |
| 11 | ENA-78 (CXCL5) | 0.15 ± 0.07 | 0.08—0.23 |
| 12 | IL-18 | 1.02 ± 1.31 | 0.14—2.92 |
| 13 | IL-1RA | 1.08 ± 0.23 | 0.83—1.28 |
| 14 | IL-29 (IFNλ1) | 0.28 ± 0.11 | 0.13—0.38 |
| 15 | IP-10 (CXCL10) | 0.70 ± 0.99 | 0.18—2.19 |
| 16 | I-TAC (CXCL11) | 1.82 ± 0.23 | 1.59—2.13 |
| 17 | MCP-1 (CCL2) | 0.14 ± 0.05 | 0.09—0.21 |
| 18 | M-CSF (CSF1) | 0.10 ± 0.04 | 0.06—0.14 |
| 19 | MIP-1β (CCL4) | 5.17 ± 2.40 | 2.78—8.51 |
| 20 | TRAIL (CD253) | 0.28 ± 0.11 | 0.18—0.44 |
| 21 | SDF1α (CXCL12) | 30.13 ± 26.73 | 10.60—60.60 |

| **b** | **Assay** | **Median LLOD**<br>**(ng/mL)** | **LLOD Range**<br>**(ng/mL)** | **Median LLOQ**<br>**(ng/mL)** | **LLOQ Range**<br>**(ng/mL)** |
| --- | --- | --- | --- | --- | --- |
| 1 | Total MPO | 0.15 ± 0.15 | 0.03—0.42 | 0.30 ± 0.30 | 0.05—0.84 |
| 2 | MPO Activity | 2.86 ± 1.98 | 0.38—6.03 | 5.72 ± 3.96 | 0.76—12.07 |

**Extended Data Table 5.** Panel of oligo-conjugated antibodies used in multi-omics scRNA-seq assays to measure surface protein markers on cells from airways and blood. A total of 89 oligo-conjugated antibody-derived tags (ADT) were used to evaluate surface protein expression of target antigens via scRNA-seq. Reagents no. 82-89 were not commercially available in the TotalSeq-C™ format (BioLegend), so custom oligo-conjugated reagents were generated in-house using purified monoclonal antibodies and commercially available 5’ Feature Barcode Antibody Conjugation - Lightning-Link^®^ kits (abcam). Titrations of all reagents were determined in-house for each lot independently prior to use. Cells were surface stained prior to encapsulation and surface marker expression of major lineage markers were used to help cluster/validate gene expression data as described in the methods.

| Antigen | Clone | Vendor | Cat. No. | Antigen | Clone | Vendor | Cat. No. |
| --- | --- | --- | --- | --- | --- | --- | --- |
| 1 CD10 | HI10a | Biolegend | 312233 | 46 CD324 | 67A4 | Biolegend | 324127 |
| 2 CD101 (BB27) | BB27 | Biolegend | 331017 | 47 CD326 (EpCAM) | 9C4 | Biolegend | 324247 |
| 3 CD103 (Integrin αE) | Ber-ACT8 | Biolegend | 350233 | 48 CD335 (NKp46) | 9.00E+02 | Biolegend | 331941 |
| 4 CD107a (LAMP-1) | H4A3 | Biolegend | 328649 | 49 CD370 (CLEC9A) | 8F9 | Biolegend | 353809 |
| 5 CD117 (c-kit) | 104D2 | Biolegend | 313243 | 50 CD38 | HIT2 | Biolegend | 303543 |
| 6 CD11b | ICRF44 | Biolegend | 301359 | 51 CD4 | RPA-T4 | Biolegend | 300567 |
| 7 CD11c | S-HCL-3 | Biolegend | 371521 | 52 CD40 | 5C3 | Biolegend | 334348 |
| 8 CD123 (IL-3Rα) | 6H6 | Biolegend | 306045 | 53 CD41 | HIP8 | Biolegend | 303739 |
| 9 CD127 (IL-7Rα) | A019D5 | Biolegend | 351356 | 54 CD44 | BJ18 | Biolegend | 338827 |
| 10 CD133 | S16016B | Biolegend | 394007 | 55 CD45 | 2D1 | Biolegend | 368545 |
| 11 CD137 | 4B4-1 | Biolegend | 309839 | 56 CD45R | RA3-6B2 | Biolegend | 103273 |
| 12 CD138 (Syndecan-1) | MI15 | Biolegend | 356539 | 57 CD45RA | HI100 | Biolegend | 304163 |
| 13 CD14 | M5E2 | Biolegend | 301859 | 58 CD45RO | UCHL1 | Biolegend | 304259 |
| 14 CD15 (SSEA-1) | W6D3 | Biolegend | 323053 | 59 CD49a | TS2/7 | Biolegend | 328319 |
| 15 CD16 (FcγRIII) | 3G8 | Biolegend | 302065 | 60 CD5 | UCHT2 | Biolegend | 300637 |
| 16 CD161 | HP-3G10 | Biolegend | 339947 | 61 CD56 (NCAM) | QA17A16 | Biolegend | 392425 |
| 17 CD163 | GHI/61 | Biolegend | 333637 | 62 CD62L | DREG-56 | Biolegend | 304851 |
| 18 CD169 | 7-239 | Biolegend | 346021 | 63 CD64 (FcγRI) | 10.1 | Biolegend | 305045 |
| 19 CD183 (CXCR3) | G025H7 | Biolegend | 353747 | 64 CD66b | 6/40c | Biolegend | 392909 |
| 20 CD184 (CXCR4) | 12G5 | Biolegend | 306533 | 65 CD69 | FN50 | Biolegend | 310951 |
| 21 CD185 (CXCR5) | J252D4 | Biolegend | 356939 | 66 CD80 | 2D10 | Biolegend | 305243 |
| 22 CD19 | HIB19 | Biolegend | 302265 | 67 CD86 | IT2.2 | Biolegend | 305447 |
| 23 CD196 (CCR6) | G034E3 | Biolegend | 353440 | 68 CD8α | RPA-T8 | Biolegend | 301071 |
| 24 CD197 (CCR7) | G043H7 | Biolegend | 353251 | 69 CD95 (Fas) | DX2 | Biolegend | 305651 |
| 25 CD1c | L161 | Biolegend | 331547 | 70 CX3CR1 | K0124E1 | Biolegend | 355705 |
| 26 CD1d | 51.1 | Biolegend | 350319 | 71 HLA-DR | L243 | Biolegend | 307663 |
| 27 CD20 | 2H7 | Biolegend | 302363 | 72 IgD | IA6-2 | Biolegend | 348245 |
| 28 CD206 (MMR) | 15-2 | Biolegend | 321147 | 73 IgG | M1310G05 | Biolegend | 410727 |
| 29 CD209 (DC-SIGN) | 9E9A8 | Biolegend | 330121 | 74 IgM | MHM-88 | Biolegend | 314547 |
| 30 CD21 | Bu32 | Biolegend | 354923 | 75 Ig light chain $\kappa$ | MHK-49 | Biolegend | 316533 |
| 31 CD23 | EBVCS-5 | Biolegend | 338525 | 76 Ig light chain $\lambda$ | MHL-38 | Biolegend | 316629 |
| 32 CD24 | ML5 | Biolegend | 311143 | 77 Siglec-8 | 7C9 | Biolegend | 347115 |
| 33 CD25 | BC96 | Biolegend | 302649 | 78 TCR Vα7.2 | 3C10 | Biolegend | 351735 |
| 34 CD269 (BCMA) | 19F2 | Biolegend | 357523 | 79 TCR $\gamma/\delta$ | B1 | Biolegend | 331231 |
| 35 CD27 | O323 | Biolegend | 302853 | 80 TIGIT (VSTM3) | A15153G | Biolegend | 372729 |
| 36 CD274 (PD-L1) | 29E.2A3 | Biolegend | 329751 | 81 XCR1 | S15046E | Biolegend | 372617 |
| 37 CD275 (ICOSL) | 2D3 | Biolegend | 309419 | **Custom oligo-conjugated in-house** |  |  |  |
| 38 CD279 (PD-1) | EH12.2H7 | Biolegend | 329963 | 82 ACE2 | 535919 | Sino Biological | MAB9332-100 |
| 39 CD28 | CD28.2 | Biolegend | 302963 | 83 CD160 | BY55 | Biolegend | 341202 |
| 40 CD29 (Integrin β1) | TS2/16 | Biolegend | 303029 | 84 CD16b | 2D2G5B9 | Sino Biological | 11046-MM01 |
| 41 CD3 | UCHT1 | Biolegend | 300479 | 85 CD178 | NOK-1 | Biolegend | 306402 |
| 42 CD307d (FcRL4) | 413D12 | Biolegend | 340213 | 86 CD34 | 581 | Biolegend | 343602 |
| 43 CD307e (FcRL5) | 509f6 | Biolegend | 340309 | 87 CD43 | CD43-10G7 | Biolegend | 343202 |
| 44 CD309 (VEGFR2) | 7D4-6 | Biolegend | 359921 | 88 CD9 | HI9a | Biolegend | 312102 |
| 45 CD32 (FcγRII) | FUN-2 | Biolegend | 303225 | 89 IgA | IS11-8E10 | Mytenyi | 130-093-073 |

**Extended Data Table 6.** Sequencing depth of independent samples. All gene expression (GEX) libraries were pooled and run in a single S4 flow cell on a NovaSeq™ 6000. All antibody-derived tag (ADT) libraries were pooled and sequenced in a single lane of a S4 flow cell on a NovaSeq™ 6000. Total reads and mean number of reads per cell for GEX and ADT are reported independently for each sample, with an average read depth of 540,000,000 reads per sample (GEX) and 79,000,000 reads per sample (ADT).

|  |  | PID | Total reads | Mean reads/cell |
| --- | --- | --- | --- | --- |
| ETA | GEX | PHA0007 | 593,835,060 | 350,345 |
|  |  | PHA0008 | 555,825,992 | 2,241,234 |
|  |  | PHA0010 | 587,153,474 | 872,442 |
|  |  | PHA0018 | 541,561,148 | 459,729 |
|  | ADT | PHA0007 | 85,070,531 | 2,101 |
|  |  | PHA0008 | 78,647,311 | 7,514 |
|  |  | PHA0010 | 46,042,783 | 2,989 |
|  |  | PHA0018 | 73,265,461 | 5,399 |
| Blood | GEX | PHA0007 | 518,672,044 | 273,129 |
|  |  | PHA0008 | 540,400,013 | 641,044 |
|  |  | PHA0010 | 592,394,112 | 543,980 |
|  |  | PHA0018 | 556,699,985 | 1,484,533 |
|  |  | PHA1005 | 541,306,254 | 158,787 |
|  |  | PHA1006 | 501,896,927 | 200,198 |
|  |  | PHA1007 | 512,999,099 | 172,902 |
|  |  | PHA1008 | 506,988,059 | 160,898 |
|  |  | PHA1009 | 527,201,752 | 130,463 |
|  |  | PHA1010 | 480,796,663 | 332,042 |
|  | ADT | PHA0007 | 74,597,277 | 2,115 |
|  |  | PHA0008 | 80,175,737 | 1,536 |
|  |  | PHA0010 | 62,927,249 | 1,067 |
|  |  | PHA0018 | 79,818,668 | 1,395 |
|  |  | PHA1005 | 77,435,258 | 1,145 |
|  |  | PHA1006 | 88,850,369 | 1,334 |
|  |  | PHA1007 | 88,329,209 | 1,169 |
|  |  | PHA1008 | 93,579,790 | 1,351 |
|  |  | PHA1009 | 91,162,676 | 1,281 |
|  |  | PHA1010 | 89,597,131 | 1,251 |

**Extended Data Table 7.** Mitochondrial gene thresholds and total ADT UMIs of individual samples. (a)Threshold for mitochondrial gene distribution (percentage) for each sample used to exclude potential dead cells in scRNA-seq data. Threshold was evaluated and set for each sample independently. (b) The distribution of total ADT UMIs was determined and recorded per each sample independently.

| **a** | **PID** | **Blood (%)** | **ETA (%)** |
| --- | --- | --- | --- |
|  | PHA0007 | 30 | 30 |
|  | PHA0008 | 20 | 25 |
|  | PHA0010 | 10 | 25 |
|  | PHA0018 | 30 | 25 |
|  | PHA1005 | 30 |  |
|  | PHA1006 | 30 |  |
|  | PHA1007 | 30 |  |
|  | PHA1008 | 20 |  |
|  | PHA1009 | 20 |  |
|  | PHA1010 | 25 |  |

| **b** | **Total ADT UMI** |  |
| --- | --- | --- |
| **PID** | **Blood** | **ETA** |
| PHA0007 | 125–500 | 125–1250 |
| PHA0008 | 125–1000 | 125–1250 |
| PHA0010 | 125–1500 | 125–750 |
| PHA0018 | 125–1500 | 125–1000 |
| PHA1005 | 125–1250 |  |
| PHA1006 | 125–1250 |  |
| PHA1007 | 125–1500 |  |
| PHA1008 | 125–1500 |  |
| PHA1009 | 125–1500 |  |
| PHA1010 | 125–1500 |  |

**Extended Data Figure 1.**
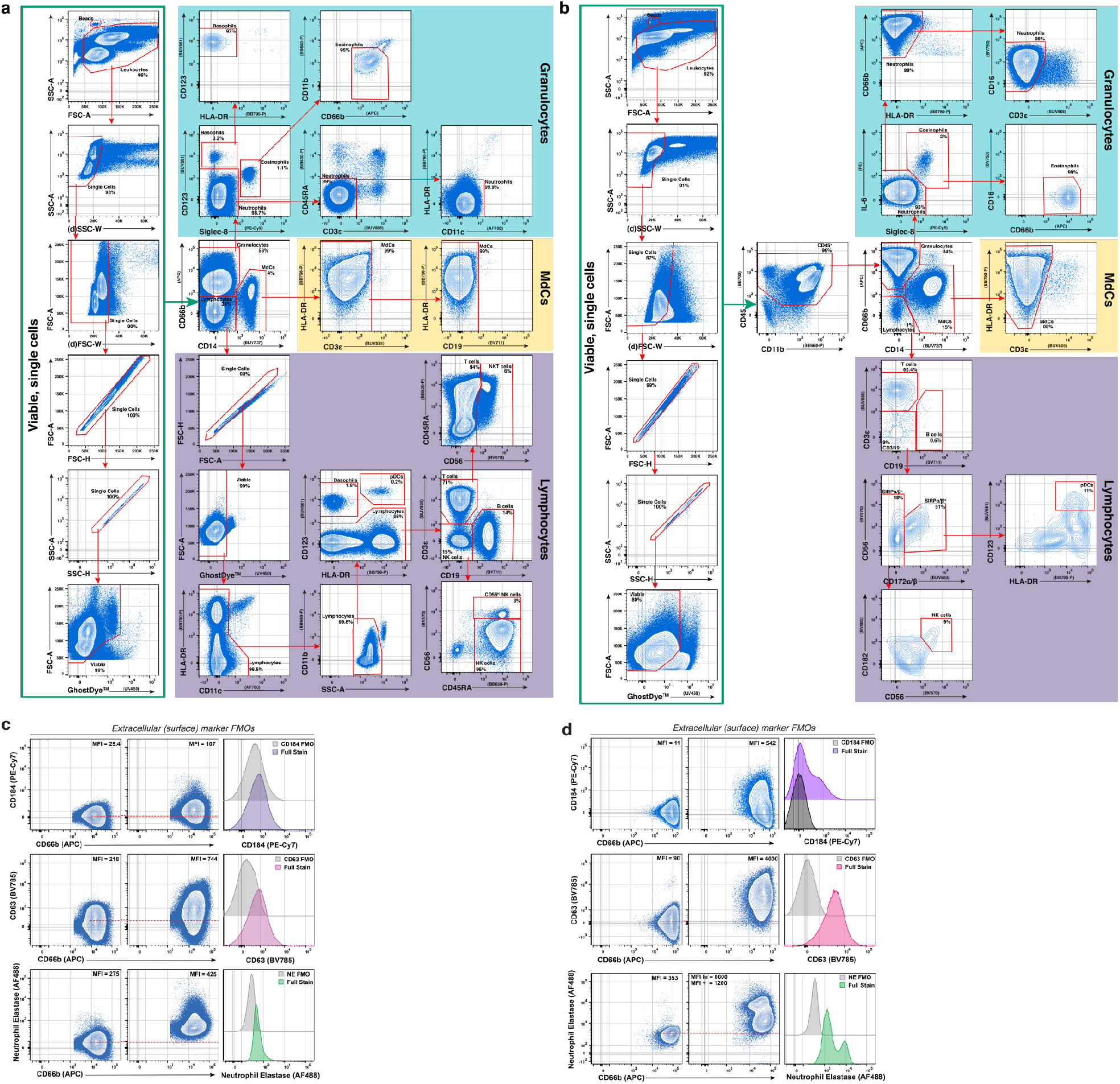
Representative gating strategy for Hi-D FACS data. Representative gating of blood (a) and ETA (b) samples. Extracellular stain FMOs for markers used to interrogate neutrophil phenotype in blood (c) and ETA (d).

**Extended Data Figure 2.**
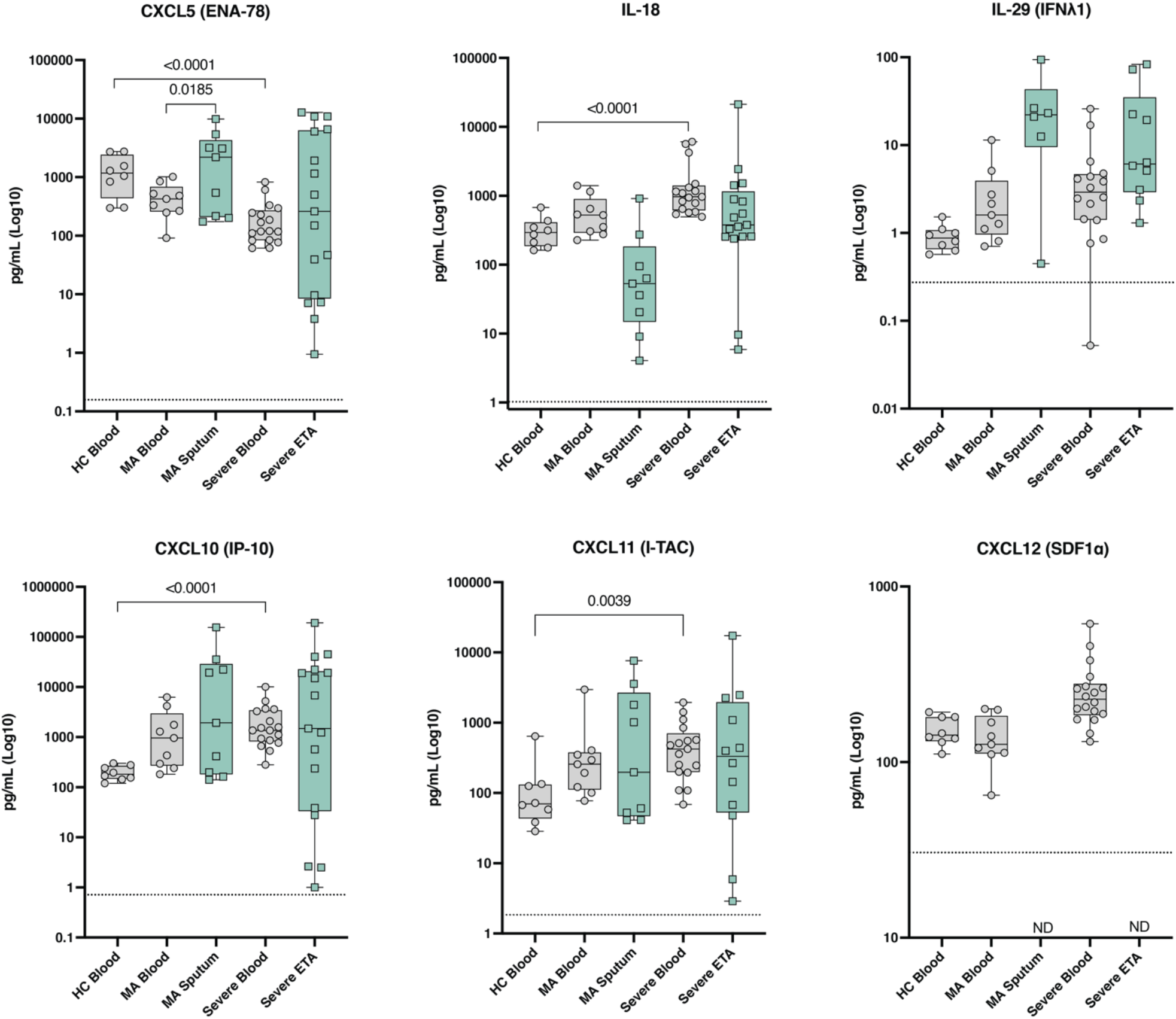
Additional cytokine assessment in blood and lungs. Concentration (pg/mL) of remaining 6 analytes interrogated by Mesoscale analyses in plasma (gray circles) and respiratory supernatant (Resp. SNT; green squares) from healthy control (HC), mild-acute (MA), and severe COVID-19 patients. Dotted line = assay limit of detection (LOD).

**Extended Data Figure 3.**
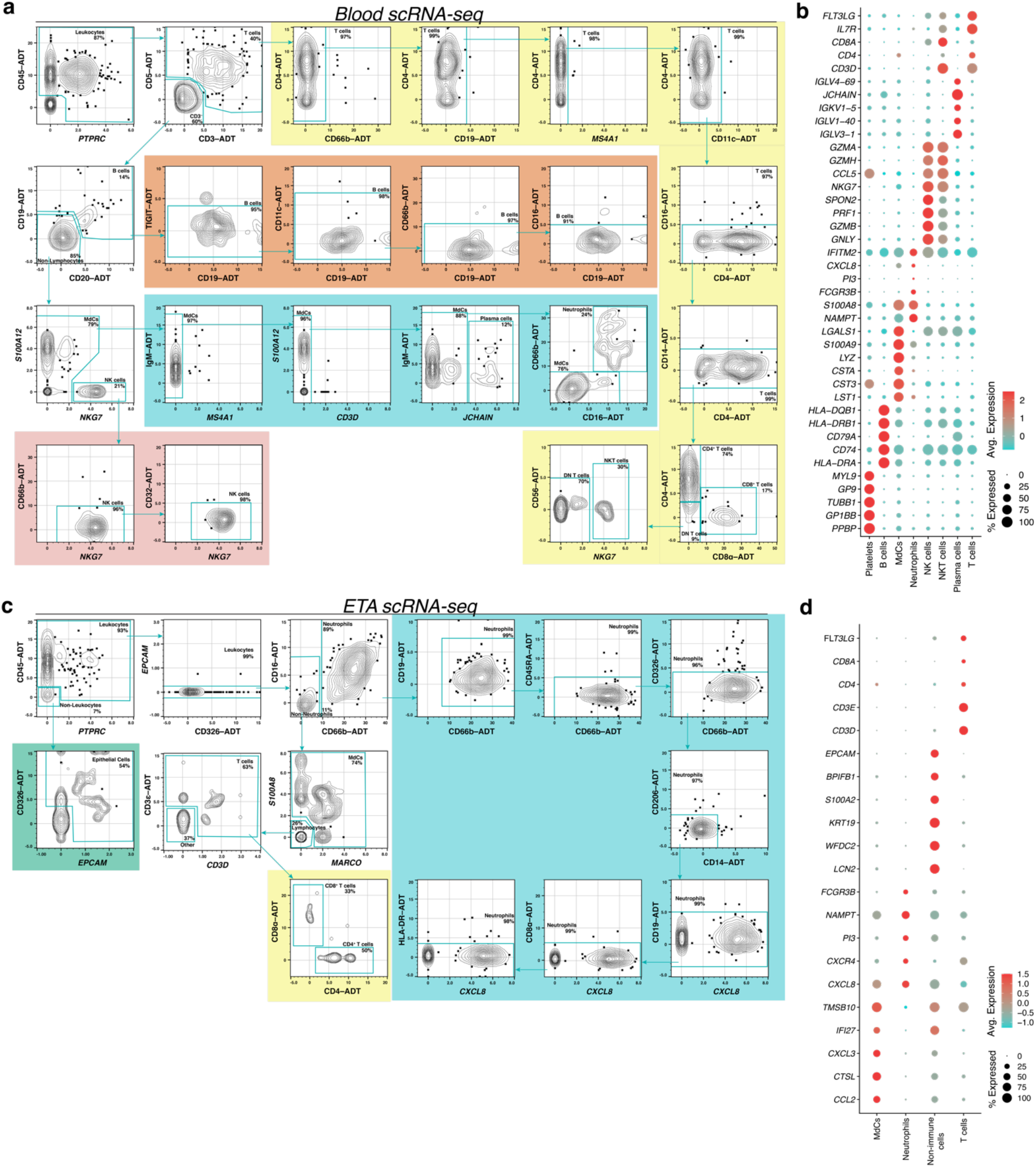
Multi-omic scRNA-seq analysis of leukocytes in the blood and lung. Gating strategy (a and c) employed to classify major lineages of immune cells by surface antibody-derived tag (ADT) and dot plots of the intersection of the top differentially expressed genes sorted by average log-fold change across cell populations (b and d) from the blood (a-b) and ETA (c-d) of severe COVID-19 patients using the SuPPER-seq pipeline previously described^30^.

**Extended Data Figure 4.**
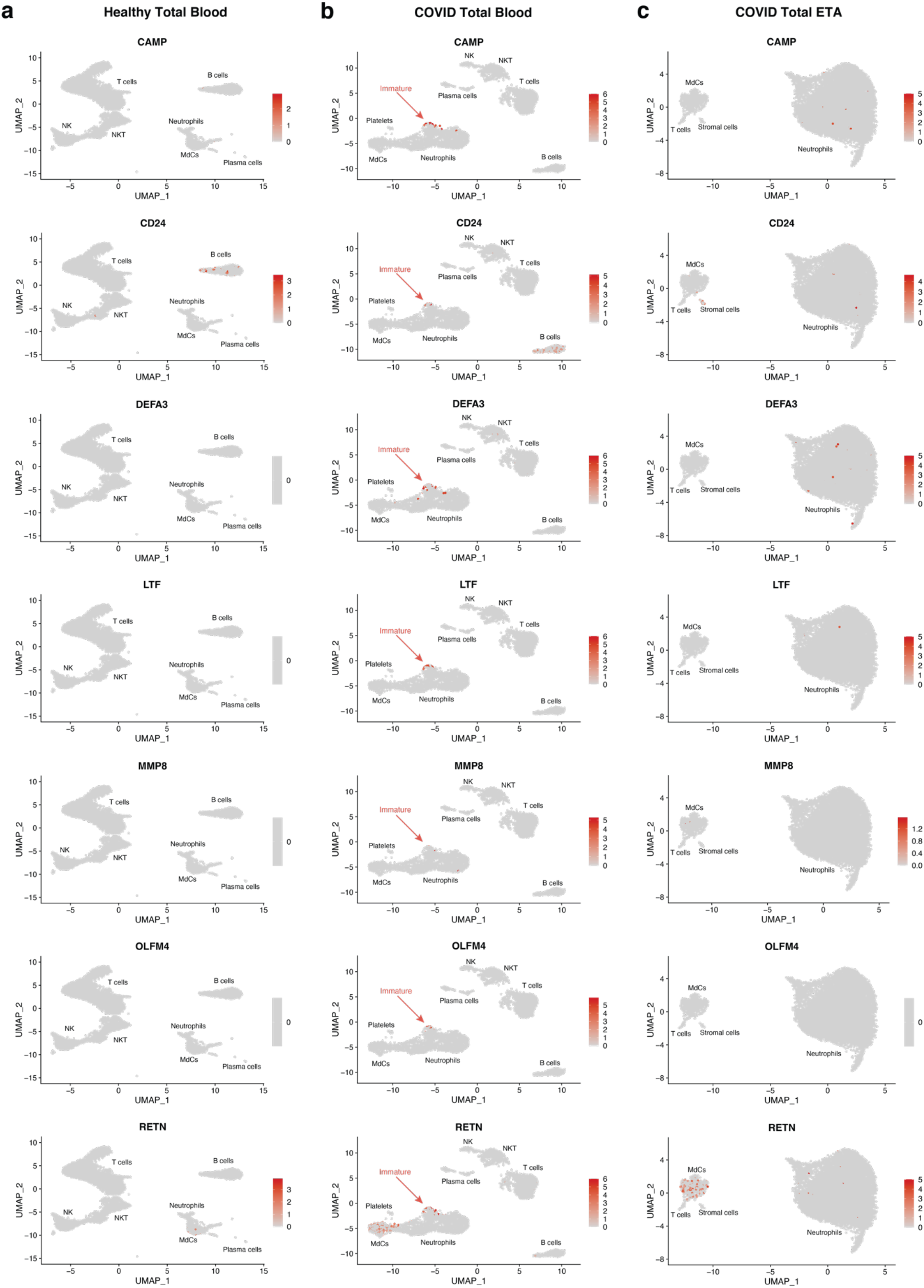
Gene signature for immature neutrophils is lacking in healthy donors and lungs of COVID-19 patients. UMAP visualizations of genes that identify immature neutrophils in the blood healthy individuals (a) and severe COVID-19 patients (b), along with cells from the lungs of severe COVID-19 patients (c).

**Extended Data Figure 5.**
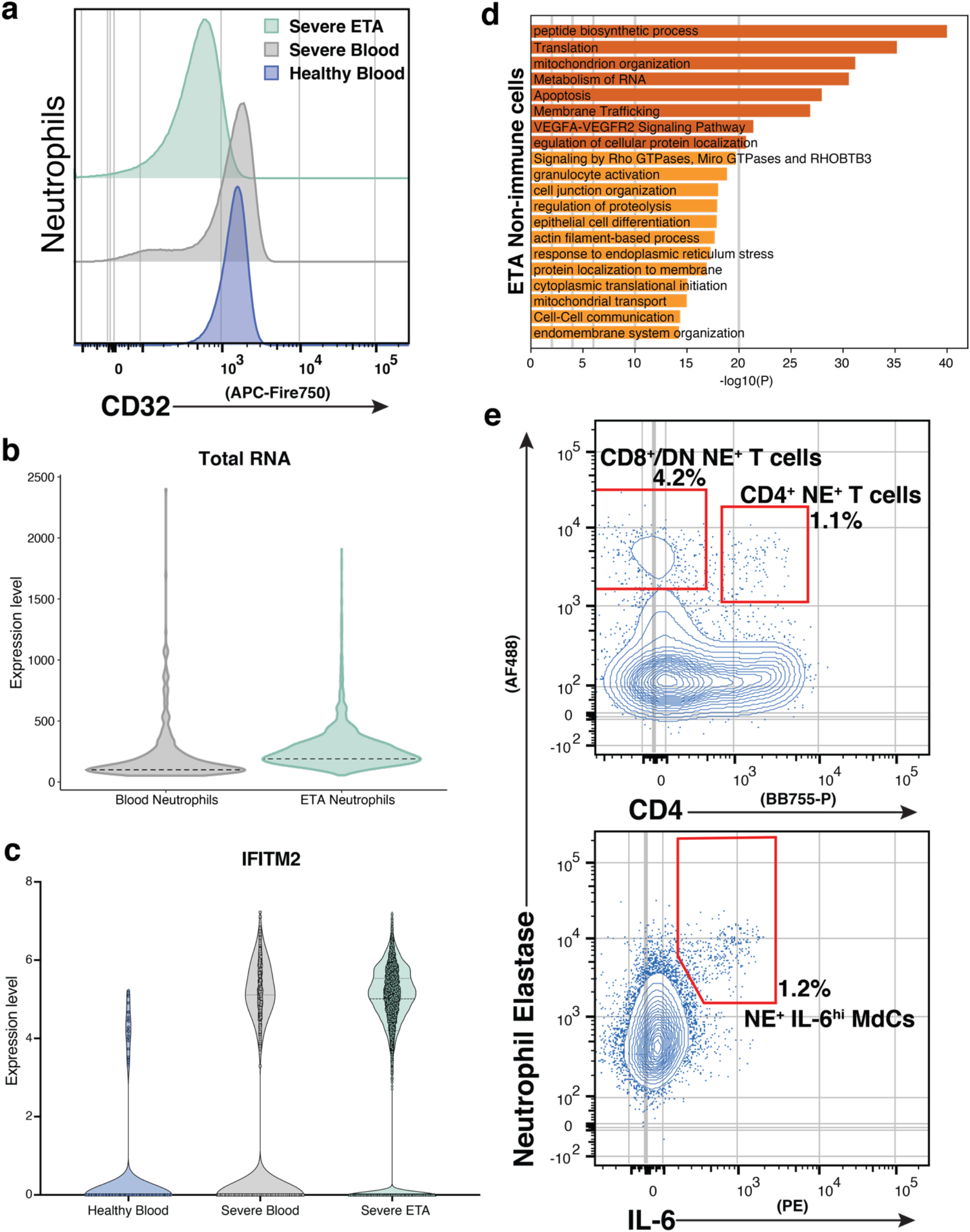
Cellular signatures during SARS-CoV-2 pathogenesis. (a) Pulmonary neutrophils have a reduction in CD32 (FcγRII) expression as compared to circulating neutrophils. (b) Neutrophils undergo transcriptional reprogramming and increase de novo transcription (total RNA) upon migrating to the lung. (c) Neutrophils in the blood and lung increase expression of *IFITM2* during SARS-CoV-2 pathogenesis. (d) Metascape^85^ pathway and process enrichment analyses reveal non-immune cells have an increased gene signature for granulocyte activation in the airways of severe COVID-19 patients. (e) Extracellular neutrophil elastase is also detected on the surface of pulmonary T cells and myeloid-derived cells (MdCs) in severe COVID-19 patients.

## References

1. Van Dyke, M.E. et al. Racial and ethnic disparities in COVID-19 incidence by age, sex, and period among persons aged< 25 years—16 US jurisdictions, January 1–December 31, 2020. Morbidity and Mortality Weekly Report 70, 382 (2021).

2. Romano, S.D. et al. Trends in Racial and Ethnic Disparities in COVID-19 Hospitalizations, by Region -United States, March-December 2020. MMWR Morb Mortal Wkly Rep 70, 560–565 (2021).

3. Lucas, C. et al. Longitudinal analyses reveal immunological misfiring in severe COVID-19. Nature 584, 463–469 (2020).

4. Merad, M. & Martin, J.C. Pathological inflammation in patients with COVID-19: a key role for monocytes and macrophages. Nature Reviews Immunology 20, 355–362 (2020).

5. Olwal, C.O. et al. Parallels in Sepsis and COVID-19 Conditions: Implications for Managing Severe COVID-19 Patients. Frontiers in immunology 12, 91 (2021).

6. Carvalho, T., Krammer, F. & Iwasaki, A. The first 12 months of COVID-19: a timeline of immunological insights. Nat Rev Immunol 21, 245–256 (2021).

7. Bost, P. et al. Deciphering the state of immune silence in fatal COVID-19 patients. Nat Commun 12, 1428 (2021).

8. Liao, M. et al. Single-cell landscape of bronchoalveolar immune cells in patients with COVID-19. Nat Med 26, 842–844 (2020).

9. Liang, W. et al. Development and Validation of a Clinical Risk Score to Predict the Occurrence of Critical Illness in Hospitalized Patients With COVID-19. JAMA Intern Med 180, 1081–1089 (2020).

10. Steinberg, K.P. et al. Evolution of bronchoalveolar cell populations in the adult respiratory distress syndrome. Am J Respir Crit Care Med 150, 113–122 (1994).

11. Barnes, B.J. et al. Targeting potential drivers of COVID-19: Neutrophil extracellular traps. J Exp Med 217 (2020).

12. Kong, M., Zhang, H., Cao, X., Mao, X. & Lu, Z. Higher level of neutrophil-to-lymphocyte is associated with severe COVID-19. Epidemiology & Infection 148 (2020).

13. Liu, Y. et al. Neutrophil-to-lymphocyte ratio as an independent risk factor for mortality in hospitalized patients with COVID-19. Journal of Infection (2020).

14. Combes, A.J. et al. Global absence and targeting of protective immune states in severe COVID-19. Nature 591, 124–130 (2021).

15. Schulte-Schrepping, J. et al. Severe COVID-19 is marked by a dysregulated myeloid cell compartment. Cell 182, 1419–1440 (2020).

16. Silvin, A. et al. Elevated Calprotectin and Abnormal Myeloid Cell Subsets Discriminate Severe from Mild COVID-19. Cell 182, 1401–1418 e1418 (2020).

17. Grant, R.A. et al. Circuits between infected macrophages and T cells in SARS-CoV-2 pneumonia. Nature 590, 635–641 (2021).

18. Wilk, A.J. et al. A single-cell atlas of the peripheral immune response in patients with severe COVID-19. Nat Med 26, 1070–1076 (2020).

19. Kreutmair, S. et al. Distinct immunological signatures discriminate severe COVID-19 from non-SARS-CoV-2-driven critical pneumonia. Immunity 54, 1578–1593 e1575 (2021).

20. Ren, X. et al. COVID-19 immune features revealed by a large-scale single-cell transcriptome atlas. Cell 184, 1895–1913 e1819 (2021).

21. Xu, G. et al. The differential immune responses to COVID-19 in peripheral and lung revealed by single-cell RNA sequencing. Cell Discov 6, 73 (2020).

22. Forrest, O.A. et al. Frontline Science: Pathological conditioning of human neutrophils recruited to the airway milieu in cystic fibrosis. J Leukoc Biol 104, 665–675 (2018).

23. Szabo, P.A. et al. Longitudinal profiling of respiratory and systemic immune responses reveals myeloid cell-driven lung inflammation in severe COVID-19. Immunity 54, 797–814 (2021).

24. Jouan, Y., Baranek, T., Si-Tahar, M., Paget, C. & Guillon, A. Lung compartmentalization of inflammatory biomarkers in COVID-19-related ARDS. Critical Care 25, 1–3 (2021).

25. Zaid, Y. et al. Chemokines and Eicosanoids Fuel the Hyperinflammation Within the Lungs of Patients with Severe COVID-19. Journal of Allergy and Clinical Immunology (2021).

26. Jouan, Y. et al. Phenotypical and functional alteration of unconventional T cells in severe COVID-19 patients. J Exp Med 217 (2020).

27. Satış, H. et al. Prognostic value of interleukin-18 and its association with other inflammatory markers and disease severity in COVID-19. Cytokine 137, 155302 (2021).

28. Yang, Y. et al. Plasma IP-10 and MCP-3 levels are highly associated with disease severity and predict the progression of COVID-19. Journal of Allergy and Clinical Immunology 146, 119–127 (2020).

29. Chandler, J.D. et al. Myeloperoxidase oxidation of methionine associates with early cystic fibrosis lung disease. Eur Respir J 52 (2018).

30. Xu, C. et al. Comprehensive multi-omics single-cell data integration reveals greater heterogeneity in the human immune system. bioRxiv, 2021.2007.2025.453651 (2021).

31. Jin, S. et al. Inference and analysis of cell-cell communication using CellChat. Nature communications 12, 1–20 (2021).

32. Bergen, V., Lange, M., Peidli, S., Wolf, F.A. & Theis, F.J. Generalizing RNA velocity to transient cell states through dynamical modeling. Nat Biotechnol 38, 1408–1414 (2020).

33. Margaroli, C. et al. Transcriptional firing represses bactericidal activity in cystic fibrosis airway neutrophils. Cell Rep Med 2, 100239 (2021).

34. Giacalone, V.D., Margaroli, C., Mall, M.A. & Tirouvanziam, R. Neutrophil Adaptations upon Recruitment to the Lung: New Concepts and Implications for Homeostasis and Disease. International Journal of Molecular Sciences 21, 851 (2020).

35. Makam, M. et al. Activation of critical, host-induced, metabolic and stress pathways marks neutrophil entry into cystic fibrosis lungs. Proc Natl Acad Sci U S A 106, 5779–5783 (2009).

36. Sercundes, M.K. et al. Targeting Neutrophils to Prevent Malaria-Associated Acute Lung Injury/Acute Respiratory Distress Syndrome in Mice. PLoS Pathog 12, e1006054 (2016).

37. Prelli Bozzo, C. et al. IFITM proteins promote SARS-CoV-2 infection and are targets for virus inhibition in vitro. Nature Communications 12, 1–13 (2021).

38. Browaeys, R., Saelens, W. & Saeys, Y. NicheNet: modeling intercellular communication by linking ligands to target genes. Nat Methods 17, 159–162 (2020).

39. Chattopadhyay, S. et al. Calmodulin binds to the cytoplasmic domain of angiotensin-converting enzyme and regulates its phosphorylation and cleavage secretion. J Biol Chem 280, 33847–33855 (2005).

40. Lambert, D.W., Clarke, N.E., Hooper, N.M. & Turner, A.J. Calmodulin interacts with angiotensin-converting enzyme-2 (ACE2) and inhibits shedding of its ectodomain. FEBS letters 582, 385–390 (2008).

41. Shimazu, R. et al. MD-2, a molecule that confers lipopolysaccharide responsiveness on Toll-like receptor 4. J Exp Med 189, 1777–1782 (1999).

42. Visintin, A., Mazzoni, A., Spitzer, J.A. & Segal, D.M. Secreted MD-2 is a large polymeric protein that efficiently confers lipopolysaccharide sensitivity to Toll-like receptor 4. Proc Natl Acad Sci U S A 98, 12156–12161 (2001).

43. Yang, H. et al. A critical cysteine is required for HMGB1 binding to Toll-like receptor 4 and activation of macrophage cytokine release. Proc Natl Acad Sci U S A 107, 11942–11947 (2010).

44. Bohn, M.K. et al. Pathophysiology of COVID-19: Mechanisms Underlying Disease Severity and Progression. Physiology (Bethesda) 35, 288–301 (2020).

45. Meizlish, M.L. et al. A neutrophil activation signature predicts critical illness and mortality in COVID-19. Blood Adv 5, 1164–1177 (2021).

46. Metzemaekers, M. et al. Kinetics of peripheral blood neutrophils in severe coronavirus disease 2019. Clin Transl Immunology 10, e1271 (2021).

47. Wauters, E. et al. Discriminating mild from critical COVID-19 by innate and adaptive immune single-cell profiling of bronchoalveolar lavages. Cell Res 31, 272–290 (2021).

48. Abedi, V. et al. Racial, economic, and health inequality and COVID-19 infection in the United States. Journal of racial and ethnic health disparities 8, 732–742 (2021).

49. Guo, Q. et al. Induction of alarmin S100A8/A9 mediates activation of aberrant neutrophils in the pathogenesis of COVID-19. Cell Host Microbe 29, 222–235 e224 (2021).

50. Hoang, T.N. et al. Baricitinib treatment resolves lower-airway macrophage inflammation and neutrophil recruitment in SARS-CoV-2-infected rhesus macaques. Cell 184, 460–475 e421 (2021).

51. Vanderheiden, A. et al. CCR2-dependent monocyte-derived cells restrict SARS-CoV-2 infection. bioRxiv (2021).

52. Krotova, K., Khodayari, N., Oshins, R., Aslanidi, G. & Brantly, M.L. Neutrophil elastase promotes macrophage cell adhesion and cytokine production through the integrin-Src kinases pathway. Sci Rep 10, 15874 (2020).

53. Towstyka, N.Y. et al. Modulation of γδ T-cell activation by neutrophil elastase. Immunology 153, 225–237 (2018).

54. Domon, H. et al. Neutrophil Elastase Subverts the Immune Response by Cleaving Toll-Like Receptors and Cytokines in Pneumococcal Pneumonia. Front Immunol 9, 732 (2018).

55. Kim, E. et al. Inhibition of elastase enhances the adjuvanticity of alum and promotes anti-SARS-CoV-2 systemic and mucosal immunity. Proc Natl Acad Sci U S A 118 (2021).

56. Delorey, T.M. et al. COVID-19 tissue atlases reveal SARS-CoV-2 pathology and cellular targets. Nature 595, 107–113 (2021).

57. Chua, R.L. et al. COVID-19 severity correlates with airway epithelium-immune cell interactions identified by single-cell analysis. Nat Biotechnol 38, 970–979 (2020).

58. Qi, F. et al. ScRNA-seq revealed the kinetic of nasopharyngeal immune responses in asymptomatic COVID-19 carriers. Cell Discov 7, 56 (2021).

59. Jenal, M. et al. The anti-apoptotic gene BCL2A1 is a novel transcriptional target of PU.1. Leukemia 24, 1073–1076 (2010).

60. Vier, J., Groth, M., Sochalska, M. & Kirschnek, S. The anti-apoptotic Bcl-2 family protein A1/Bfl-1 regulates neutrophil survival and homeostasis and is controlled via PI3K and JAK/STAT signaling. Cell Death Dis 7, e2103 (2016).

61. Woehrl, B. et al. CXCL16 contributes to neutrophil recruitment to cerebrospinal fluid in pneumococcal meningitis. J Infect Dis 202, 1389–1396 (2010).

62. Zhang, L. et al. Chemokine CXCL16 regulates neutrophil and macrophage infiltration into injured muscle, promoting muscle regeneration. Am J Pathol 175, 2518–2527 (2009).

63. Besteman, S.B. et al. Transcriptome of airway neutrophils reveals an interferon response in life-threatening respiratory syncytial virus infection. Clinical Immunology 220, 108593 (2020).

64. Ballesteros, I. et al. Co-option of Neutrophil Fates by Tissue Environments. Cell 183, 1282–1297 e1218 (2020).

65. Yamada, M. et al. The increase in surface CXCR4 expression on lung extravascular neutrophils and its effects on neutrophils during endotoxin-induced lung injury. Cell Mol Immunol 8, 305–314 (2011).

66. Bernhagen, J. et al. MIF is a noncognate ligand of CXC chemokine receptors in inflammatory and atherogenic cell recruitment. Nature medicine 13, 587–596 (2007).

67. Pawig, L., Klasen, C., Weber, C., Bernhagen, J. & Noels, H. Diversity and Inter-Connections in the CXCR4 Chemokine Receptor/Ligand Family: Molecular Perspectives. Front Immunol 6, 429 (2015).

68. Rodrigues, D.A.S. et al. CXCR4 and MIF are required for neutrophil extracellular trap release triggered by Plasmodium-infected erythrocytes. PLoS pathogens 16, e1008230 (2020).

69. Adrover, J.M. et al. A neutrophil timer coordinates immune defense and vascular protection. Immunity 50, 390–402 (2019).

70. Casanova-Acebes, M. et al. Rhythmic modulation of the hematopoietic niche through neutrophil clearance. Cell 153, 1025–1035 (2013).

71. Ng, L.G., Ostuni, R. & Hidalgo, A. Heterogeneity of neutrophils. Nature Reviews Immunology 19, 255–265 (2019).

72. Neidleman, J. et al. Distinctive features of SARS-CoV-2-specific T cells predict recovery from severe COVID-19. medRxiv (2021).

73. Group, R.C. Dexamethasone in hospitalized patients with Covid-19. New England Journal of Medicine 384, 693–704 (2021).

74. Ronchetti, S., Ricci, E., Migliorati, G., Gentili, M. & Riccardi, C. How glucocorticoids affect the neutrophil life. International journal of molecular sciences 19, 4090 (2018).

75. Mullard, A. Anti-IL-6Rs falter in COVID-19. Nat Rev Drug Discov 19, 577 (2020).

76. Eddins, D.J. et al. Inactivation of SARS Coronavirus 2 and COVID-19 patient samples for contemporary immunology studies. bioRxiv (2021).

77. Beigel, J.H. et al. Remdesivir for the Treatment of Covid-19 -Final Report. N Engl J Med 383, 1813–1826 (2020).

78. Orlov, M. et al. Endotracheal aspirates contain a limited number of lower respiratory tract immune cells. Critical Care 25, 1–3 (2021).

79. Seren, S. et al. Proteinase release from activated neutrophils in mechanically ventilated patients with non-COVID-19 and COVID-19 pneumonia. Eur Respir J 57 (2021).

80. Dobin, A. et al. STAR: ultrafast universal RNA-seq aligner. Bioinformatics 29, 15–21 (2013).

81. Hao, Y. et al. Integrated analysis of multimodal single-cell data. bioRxiv (2020).

82. Babcock, B.R., Kosters, A., Yang, J., White, M.L. & Ghosn, E. Data Matrix Normalization and Merging Strategies Minimize Batch-specific Systemic Variation in scRNA-Seq Data. bioRxiv (2021).

83. Mulè, M.P., Martins, A.J. & Tsang, J.S. Normalizing and denoising protein expression data from droplet-based single cell profiling. bioRxiv (2020).

84. Petukhov, V. et al. dropEst: pipeline for accurate estimation of molecular counts in droplet-based single-cell RNA-seq experiments. Genome Biol 19, 78 (2018).

85. Zhou, Y. et al. Metascape provides a biologist-oriented resource for the analysis of systems-level datasets. Nat Commun 10, 1523 (2019).

86. Lu, R. et al. Genomic characterisation and epidemiology of 2019 novel coronavirus: implications for virus origins and receptor binding. Lancet 395, 565–574 (2020).

